# Repurposing native non-homologous end joining for multicopy random integration in *Wickerhamomyces ciferrii*

**DOI:** 10.64898/2026.05.17.725789

**Authors:** Seong-Rae Lee, Yubeen Seo, Pyung Cheon Lee

**Author notes:** Corresponding Author: Prof. Pyung Cheon Lee, Department of Molecular Science and Technology and Advanced College of Bio-convergence Engineering, Ajou University, Woncheon-dong, Yeongtong-gu, Suwon 16499, South Korea.

## Abstract

*Wickerhamomyces ciferrii* is a non-model diploid yeast that naturally produces tetraacetyl phytosphingosine (TAPS), a sphingoid base used in cosmetic and dermatological applications. However, its strong preference for non-homologous end joining (NHEJ) over homologous recombination (HR) limits conventional genome editing, while disruption of *LIG4*, a core NHEJ gene, compromises cellular fitness. Here, we repurposed native NHEJ activity to develop a homology-independent multicopy genome integration platform for *W. ciferrii*. The platform combines three optimized donor-design features: telomeric end-shielding with two tandem copies of an 11 bp repeat to improve linear donor persistence, a defective *URA5* auxotrophic marker to enrich multicopy integrants, and 5′-phosphorylated donor termini to enhance transformant recovery and integration output. These features were consolidated into the platform vector pTdmVU5. As a metabolic engineering demonstration, multicopy integration of *LCB1* and *LCB2*, encoding the two subunits of serine palmitoyltransferase, increased TAPS titer by 2.7-fold. This work converts the native NHEJ bias of *W. ciferrii* from a barrier to precise genome editing into a practical tool for pathway amplification and establishes a framework for engineering NHEJ-dominant non-model yeasts.

## Introduction

Microbial cell factories are central to sustainable biomanufacturing because they enable the production of chemicals, pharmaceuticals, nutraceuticals, and cosmetic ingredients from renewable feedstocks. Among eukaryotic hosts, *Saccharomyces cerevisiae* has long served as a dominant model yeast owing to its well-characterized genetics, robust homologous recombination (HR), and extensive synthetic biology toolbox. However, no single chassis is optimal for all products. The metabolic range, stress tolerance, secretion capacity, and endogenous precursor availability of *S. cerevisiae* can limit its suitability for certain industrial targets, motivating the development of non-model yeasts with specialized physiological and metabolic traits.

Non-model yeasts are particularly attractive when their native metabolism is already aligned with the desired product class.^1,2^ In model hosts, construction of heterologous pathways often requires iterative cycles of enzyme screening, pathway reconstruction, genome integration, regulatory optimization, and flux balancing. In contrast, naturally predisposed yeasts can provide endogenous precursor supply, product-related metabolic enzymes, and tolerance mechanisms that reduce the engineering burden. Several such hosts have been developed for specialized applications. *Yarrowia lipolytica* is widely used for lipid-derived biochemicals because of its strong flux toward acetyl-CoA and lipid metabolism.^3–8^ *Komagataella phaffii* (also referred to as *Pichia pastoris*) is a preferred platform for recombinant protein secretion under strong inducible promoters.^9,10^ *Rhodotorula toruloides* efficiently converts lignocellulosic sugars into acetyl-CoA–derived products.^11–14^ *Pichia kudriavzevii* tolerates low pH, elevated temperature, and osmotic stress, making it useful for organic acid production.^15–19^ These examples illustrate the value of expanding yeast synthetic biology beyond conventional model systems.

Among emerging non-model yeasts, *Wickerhamomyces ciferrii* is a promising chassis for sphingolipid-related biomanufacturing. This yeast naturally produces and secretes tetraacetyl phytosphingosine (TAPS), a high-value sphingoid base used in cosmetic and dermatological products.^20,21^ It also grows efficiently under defined conditions, with reported specific growth rates of 0.27–0.64 h^-1^ in minimal medium,^22^ and has demonstrated robust fermentation performance. In addition, the α-mating factor secretion signal derived from *W. ciferrii* outperformed the corresponding *S. cerevisiae* signal when tested in *K. phaffii*, suggesting that some secretion-associated elements from *W. ciferrii* may function efficiently across yeast species.^23^ Together with its native sphingolipid metabolism, these features make *W. ciferrii* an attractive platform for sustainable production of TAPS and related bioactive molecules.

Despite this promise, genome engineering in *W. ciferrii* remains substantially less developed than in model yeasts. Previous metabolic engineering strategies improved TAPS production by overexpressing sphingolipid biosynthetic genes such as *LCB1, LCB2,* and *SYR2,* or by disrupting competing reactions such as *LCB4*-mediated phosphorylation (**Figure S1**).^24,25^ However, many of these approaches relied on HR-based manipulation in strains presumed to be haploid. Subsequent genomic analyses revealed that industrial *W. ciferrii* production strains are diploid and exhibit considerable allelic heterogeneity, making biallelic gene disruption more difficult. Moreover, the low genomic G + C content of *W. ciferrii*, approximately 30.4%,^26^ creates a distinct codon usage pattern that can influence heterologous expression depending on codon adaptation index (CAI) values.^27^ In addition to these genomic constraints, *W. ciferrii* exhibits a strong intrinsic bias toward non-homologous end joining (NHEJ), which limits the efficiency of homology-directed genome editing.^28^

NHEJ-dominant DNA repair is commonly viewed as a barrier to precise genome engineering in non-model yeasts.^29^ A frequent strategy to improve HR-mediated editing is to disrupt core NHEJ components such as DNA ligase IV. However, permanent suppression of NHEJ can impose fitness costs, particularly in industrial strains where genome stability and production performance are critical. More importantly, the same repair pathway that hinders precise HR-based editing can be repurposed for homology-independent random integration. NHEJ-mediated integration has been used to promote multicopy gene insertion and pathway amplification in several non-model yeasts, including *Y. lipolytica*, *Kluyveromyces marxianus*, and *Scheffersomyces stipitis*.^30–40^ These studies suggest that NHEJ can be transformed from a genome-editing obstacle into a useful mechanism for rapid strain construction, especially in hosts where targeted editing is inefficient or physiologically burdensome.

For *W. ciferrii*, however, a practical NHEJ-based multicopy integration platform has not been established. Recent advances in functional markers, promoters, episomal vectors, and fluorescent reporters have begun to expand the genetic toolbox for this yeast,^27^ but several barriers still limit large-scale pathway amplification. First, linear donor DNA can be degraded by endogenous nucleases before genomic capture. Second, conventional selection markers primarily identify successful transformants and do not necessarily enrich high-copy integrants. Third, donor DNA end chemistry and structure may affect recognition and ligation by the NHEJ machinery. Addressing these design constraints is essential for converting the native NHEJ bias of *W. ciferrii* into a robust synthetic biology tool.

Here, we developed a homology-independent multicopy genome integration platform that harnesses native NHEJ activity in diploid *W. ciferrii* (**Figure 1**). Rather than disrupting NHEJ to force HR-mediated editing, we optimized linear donor architecture to improve random integration efficiency and copy-number enrichment. The platform incorporates three design principles: telomeric end-shielding to enhance donor DNA persistence, defective *URA5* marker selection to enrich multicopy integrants, and 5′-phosphorylated donor termini to improve transformant recovery and reporter output. These elements were consolidated into the platform vector pTdmVU5 and validated by multicopy integration of *LCB1* and *LCB2*, which encode the two subunits of serine palmitoyltransferase. The resulting strains showed increased gene dosage, elevated transcript abundance, and enhanced TAPS production. This work establishes a practical genome engineering framework for pathway amplification in *W. ciferrii* and provides a general strategy for exploiting NHEJ-dominant DNA repair in diploid non-model yeasts.

**Figure 1.**
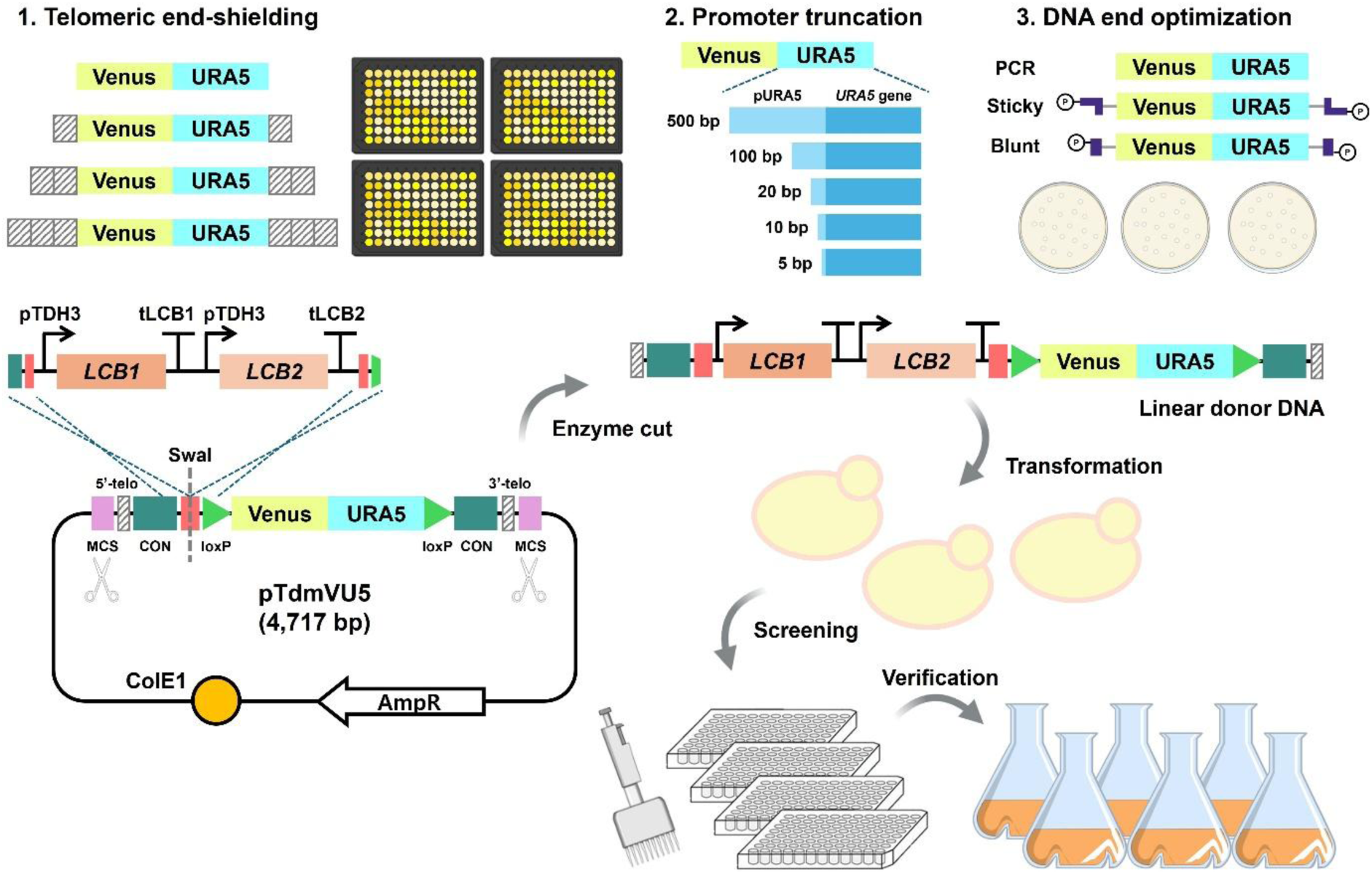
Design and workflow of an NHEJ-based multicopy genome integration platform in *W. ciferrii*. The platform exploits the native non-homologous end joining (NHEJ) bias of *W. ciferrii* for homology-independent genome integration. Three donor-design parameters were optimized: telomeric end-shielding to improve linear donor stability, *URA5* promoter truncation to generate a defective auxotrophic marker for multicopy enrichment, and donor-end configuration to enhance NHEJ-compatible insertion. These elements were integrated into the platform vector pTdmVU5. Pathway cassettes, exemplified by *LCB1* and *LCB2*, are assembled into pTdmVU5, released by restriction enzyme digestion as linear donor DNA, and transformed into Δ*URA5* cells. Transformants are screened by Venus fluorescence in 96-deep-well plates and validated by flask-scale cultivation and production analysis.

## Results and Discussion

### Native NHEJ dominance provides a basis for homology-independent genome integration in *W. ciferrii*

Previous genome engineering in *W. ciferrii* has primarily sought to improve homologous recombination (HR)-mediated targeting by disrupting *LIG4*, which encodes DNA ligase IV, a core enzyme in the non-homologous end joining (NHEJ) pathway.^24,28^ In haploid *W. ciferrii*, disruption of a 500 bp internal fragment of *LIG4* by single-crossover recombination yielded only 4–5% targeted integration, whereas double-crossover strategies using 400–500 bp homology arms achieved less than 1% efficiency.^28^ These findings indicate that *W. ciferrii* has intrinsically low HR activity. Consistently, deletion of *LIG4* increased targeting efficiency to approximately 87%, demonstrating that HR-based genome editing becomes feasible when NHEJ is genetically suppressed.^28^

To directly quantify DNA repair pathway preference in *W. ciferrii*, we adapted a split-Venus reporter assay originally developed by Ploessl et al.^38^ The Venus coding sequence was divided into two fragments sharing 50 bp terminal homology. Co-transformation of these fragments allows HR-mediated repair to be quantified by fluorescence restoration: HR reconstructs a functional Venus gene, whereas NHEJ-mediated ligation or indel formation generally fails to restore the correct reading frame and produces non-fluorescent colonies (**Figure 2A**). As expected, nearly all *S. cerevisiae* transformants exhibited Venus fluorescence, confirming its strong HR bias.^38^ In contrast, only approximately 6% of *W. ciferrii* transformants were fluorescent (**Figure 2B**). This HR frequency was lower than previously reported values for *Y. lipolytica* approximately 20% and for *K. marxianus* or *S. stipitis* approximately 10%,^38^ indicating that *W. ciferrii* is an exceptionally NHEJ-dominant yeast.

**Figure 2.**
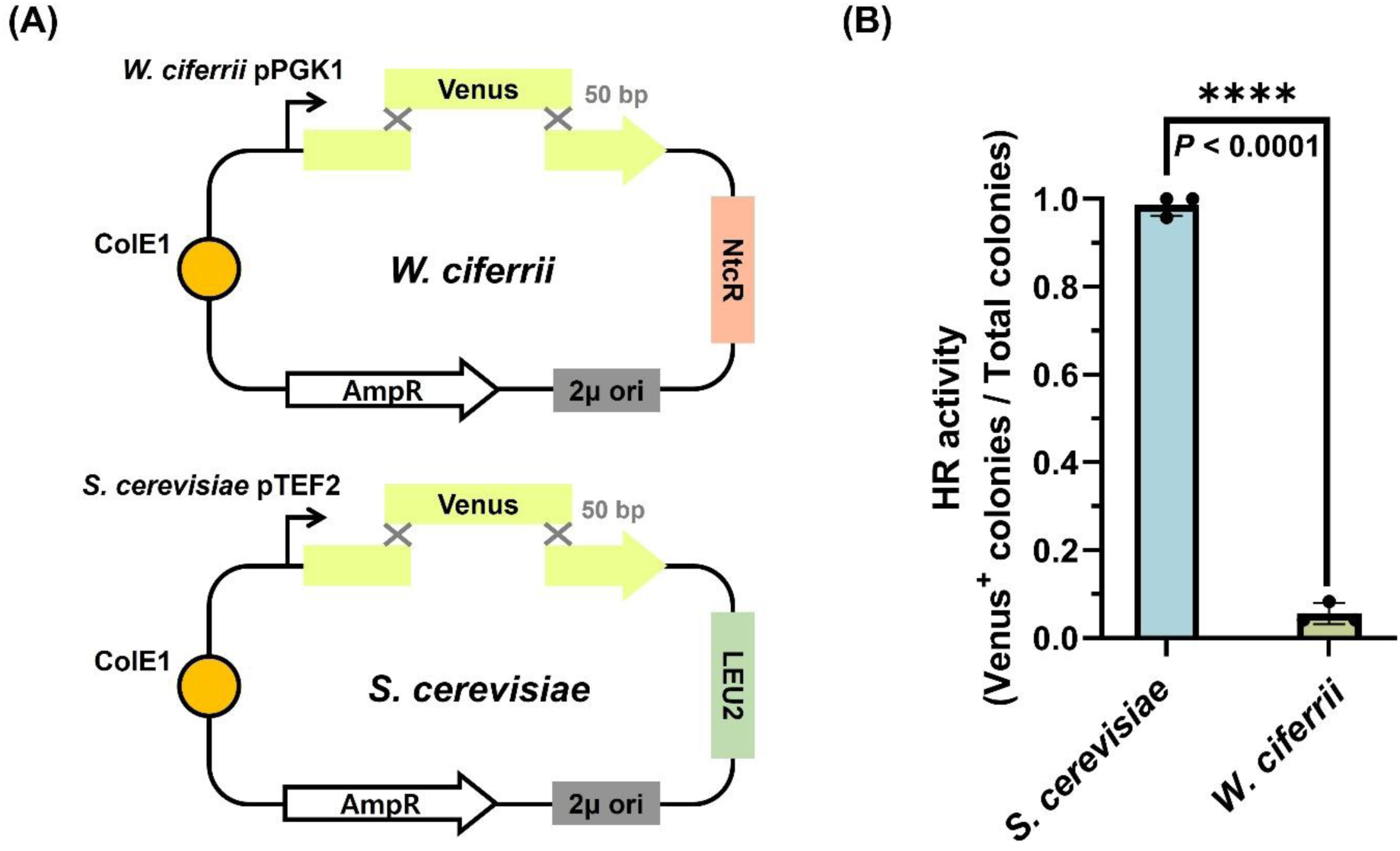
Quantitative assessment of DNA repair pathway preference in *W. ciferrii* using a split-Venus reporter assay. **(A)** Schematic of the split-Venus assay used to distinguish homologous recombination (HR)-mediated repair from non-homologous end joining (NHEJ)-mediated repair. The Venus coding sequence was divided into two fragments sharing a 50 bp terminal overlap. HR restores a functional Venus gene and generates fluorescent colonies, whereas NHEJ-mediated ligation or indel formation generally fails to reconstitute Venus fluorescence. The assay was performed in *S. cerevisiae* and *W. ciferrii*. **(B)** HR activity, calculated as the percentage of Venus-positive colonies among total transformants. *S. cerevisiae* showed near-complete fluorescence restoration, whereas *W. ciferrii* exhibited markedly low HR activity, indicating a strong NHEJ bias. Data represent the mean ± SD from three independent experiments, with 32 transformants analyzed per replicate. Statistical significance was determined using an unpaired *t*-test with Welch’s correction (\*\*\*\**p* < 0.0001).

Although NHEJ disruption can improve HR-mediated editing, this strategy is less suitable for industrial *W. ciferrii* production strains, which are diploid rather than haploid. To evaluate whether *LIG4* knockout is practical in a diploid production background, we constructed two disruption vectors carrying nourseothricin and geneticin resistance markers, respectively, and sequentially targeted both *LIG4* alleles (**Figure S2A**). Biallelic disruption was inefficient, yielding positive transformants at a frequency of approximately 1 in 40 colonies for each allele. Cre recombinase-mediated marker recycling was also highly inefficient, with a recovery frequency below 1%, consistent with previous reports, and marker-free clones could not be readily obtained. Most importantly, the resulting Δ*LIG4* strain showed reduced cellular fitness. The maximum OD_600_ decreased by approximately 35% compared with the wild-type control carrying empty episomal plasmids under antibiotic selection (**Figure S2B**), and tetraacetyl phytosphingosine (TAPS) titer decreased by approximately 13% (**Figure S2C**). These results indicate that permanent *LIG4* disruption imposes a physiological burden and is not an optimal strategy for engineering diploid *W. ciferrii* production strains.

This fitness cost is consistent with observations in other yeasts, where disruption of NHEJ-associated genes can impair cellular robustness. In *S. cerevisiae*, deletion of KU homologs *HDF1* and *HDF2*, which encode the DNA end-binding complex that initiates NHEJ, causes temperature-sensitive growth arrest and increased DNA damage sensitivity under stress conditions.^41^ Similarly, *KU70* disruption in *K. phaffii* reduced the specific growth rate and increased UV sensitivity,^42^ whereas *KU80* disruption in *O. polymorpha* caused retarded growth and reduced CFU/OD_600_.^43^ By contrast, transient repression of *KU70* and *KU80* by CRISPR interference in *Y. lipolytica* improved HR efficiency without detectable growth defects, highlighting the advantage of avoiding permanent NHEJ disruption when host fitness is critical.^44^

We next asked whether the pronounced NHEJ bias of *W. ciferrii* could be repurposed for homology-independent genome integration. Because we previously established an autonomously replicating episomal plasmid system in *W. ciferrii*,^27^ we compared the transformation behavior of circular plasmids and linear donor DNA. Donor topology had a marked effect on transformant recovery: blunt-ended linear DNA yielded approximately 2,500 CFU/μg DNA, whereas circular plasmids yielded only approximately 150 CFU/μg DNA, corresponding to approximately 17-fold higher recovery of transformants. In addition, episomal reporter plasmids produced heterogeneous fluorescence among colonies, whereas PCR-amplified linear antibiotic-resistance cassettes generated stable transformants under selection. These results are consistent with efficient capture and maintenance of linear donor DNA in *W. ciferrii*, likely through random genomic integration mediated by its native NHEJ machinery. Thus, rather than serving solely as a barrier to HR-based editing, endogenous NHEJ provides a practical entry point for homology-independent genome integration.

Together, these findings establish *W. ciferrii* as a strongly NHEJ-dominant host in which permanent suppression of NHEJ compromises the fitness and production capacity of diploid strains. We therefore sought to repurpose native NHEJ activity as a mechanism for homology-independent multicopy genome integration. This strategy led to the design of an integration platform that combines linear donor stabilization, multicopy-selective marker pressure, and donor-end optimization to enhance NHEJ-mediated genome insertion.

### Δ*URA5* auxotrophic strain and VU5 reporter cassette enable fluorescence-based monitoring of random integration

NHEJ-mediated random integration enables homology-independent insertion of linear donor DNA and can facilitate multicopy gene introduction without designing locus-specific homology arms. To adapt this strategy to *W. ciferrii*, we first established a uracil auxotrophic recipient strain and a fluorescence-based reporter cassette that couples auxotrophic selection and with quantitative reporter output.

In *W. ciferrii*, available selection strategies include antibiotic resistance markers and uracil auxotrophy-based selection and counterselection. Uracil auxotrophy is particularly useful because it enables positive selection on uracil dropout medium and counterselection on 5-fluoroorotic acid (5-FOA), providing a basis for marker recycling. To generate a uracil auxotrophic host, diploid *W. ciferrii* was serially cultured under increasing 5-FOA concentrations from 0.1 to 2.5 g/L. The resulting isolate failed to grow on SD-Ura agar, whereas wild-type cells grew normally, confirming uracil auxotrophy. Conversely, wild-type cells did not form colonies on 5-FOA agar, consistent with an intact uracil biosynthetic pathway.

Because uracil auxotrophy in yeasts is more commonly associated with loss of *URA3* function, we first examined the *URA3* locus. Genomic PCR showed that *URA3* (Gene ID: BN7_2071; RefSeq: NW_011887692.1), which encodes orotidine 5′-phosphate decarboxylase, remained intact in the auxotrophic isolate. We therefore examined *URA5*, which encodes orotate phosphoribosyltransferase and has been used as a functional selection marker in several yeasts, including *K. phaffii* and *K. lactis*.^45,46^ Although *URA5* disruption can produce leaky auxotrophic phenotypes in yeasts containing redundant orotate phosphoribosyltransferase activity, such as *S. cerevisiae* and *Y. lipolytica*,^47,48^ genomic PCR failed to amplify *URA5* (Gene ID: BN7_4897; RefSeq: NW_011887552.1) in the selected *W. ciferrii* isolate, consistent with disruption or loss of this locus. Complementation with a PCR-amplified *URA5* cassette containing 500 bp upstream and 140 bp downstream sequences restored growth on SD-Ura medium and abolished 5-FOA resistance. These results confirmed that loss of *URA5*, rather than *URA3*, was responsible for the uracil auxotrophic phenotype.

We next constructed a dual-function VU5 reporter cassette to monitor NHEJ-mediated random integration. In this cassette, Venus fluorescent protein is driven by the endogenous *PGK1* promoter and *ENO1* terminator and linked to a functional *URA5* marker. This design enables selection of *URA5*-complemented transformants while allowing Venus fluorescence to serve as a quantitative proxy for reporter expression, donor integrity, and potential copy-number variation. The VU cassette was cloned into the pUCM backbone to generate pVU5, PCR-amplified as a 2,865 bp linear fragment, and transformed into Δ*URA5* competent cells (**Figure 3A**).

**Figure 3.**
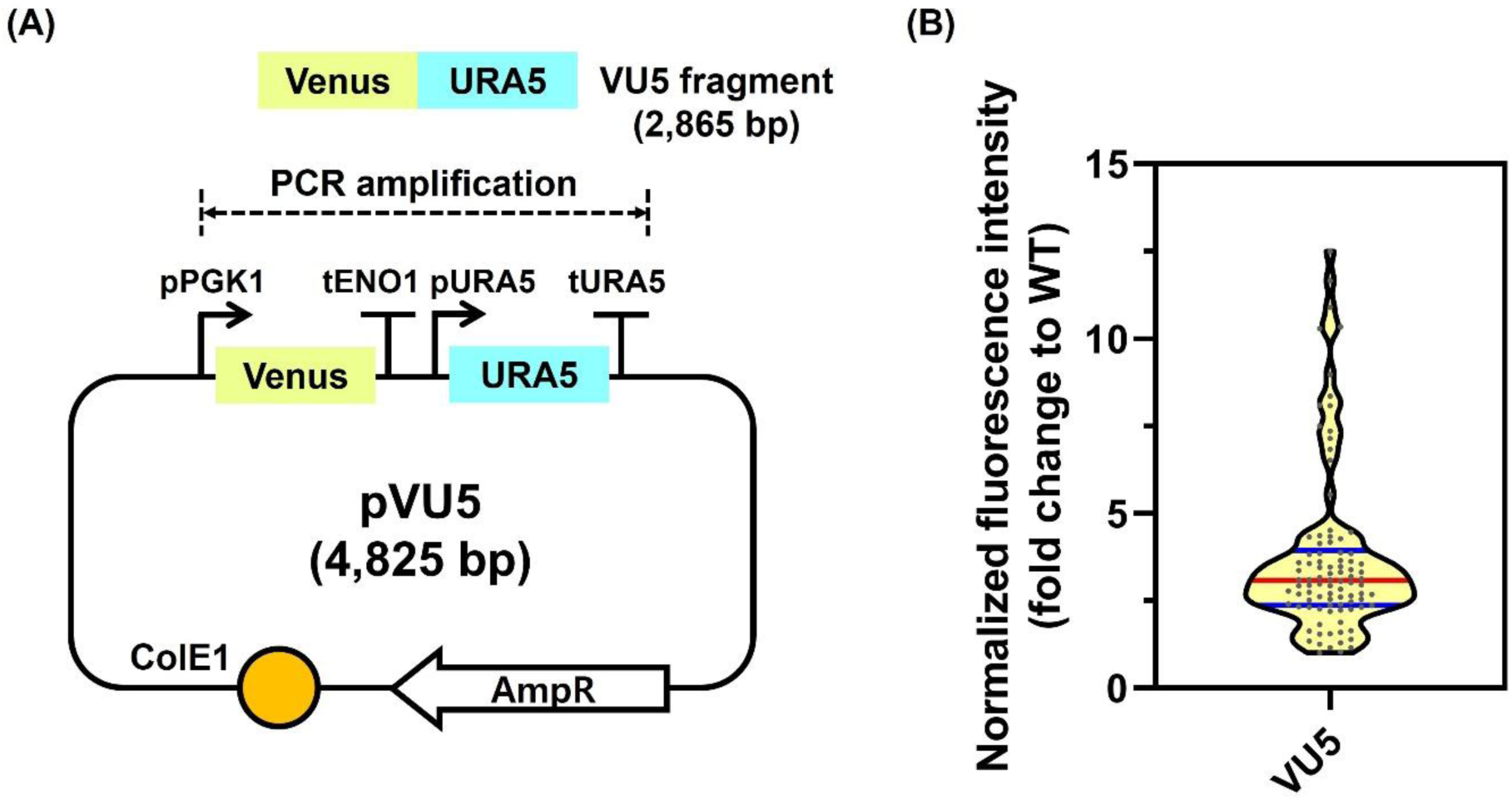
VU5 reporter system for monitoring random integration in Δ*URA5 W. ciferrii*. **(A)** Design of the pVU5 plasmid and PCR-amplified VU5 linear donor fragment. Venus is expressed from the endogenous *PGK1* promoter and *ENO1* terminator, and linked to a functional *URA5* selection marker under its native promoter and terminator. The 2,865 bp VU5 fragment was amplified from the pUCM-based pVU5 plasmid for transformation into Δ*URA5* cells. **(B)** Normalized Venus fluorescence distribution of VU5 transformants after 48 h cultivation in YMglSC medium. Fluorescence was normalized to OD_600_ and expressed relative to the non-fluorescent control. The broad distribution of reporter output suggests heterogeneous integration outcomes, including differences in genomic position, donor integrity, or copy number. Red and blue lines indicate the median and interquartile range, respectively.

To evaluate reporter output, 93 transformants and three non-fluorescent controls were cultured in 96-deep-well plates containing 1 mL of YMglSC medium at 25 °C and 800 rpm for 48 h. Venus fluorescence was normalized to OD_600_ and then to the non-fluorescent control. After outlier removal, normalized fluorescence values displayed a broad distribution, ranging from approximately 1- to 12.5-fold relative to the control (**Figure 3B**). This heterogeneous fluorescence profile is consistent with random integration outcomes that vary in reporter expression, donor integrity, genomic context, or copy number. Together, these results establish the Δ*URA5* strain and VU5 cassette as a practical selection and screening system for monitoring homology-independent random integration in *W. ciferrii*.

### Telomeric end-shielding improves linear donor persistence and random integration output

The broad fluorescence distribution of VU5 transformants suggested that random integration in *W. ciferrii* may be influenced not only by donor copy number and genomic position, but also by donor integrity during transformation. Because linear DNA ends are vulnerable to exonucleolytic processing before genomic capture, we asked whether short terminal repeat sequences could protect the internal donor cassette and improve NHEJ-mediated integration output. Telomeres function as protective chromosome-end structures that prevent degradation, end-to-end fusion, and inappropriate DNA damage responses.^49,50^ Moreover, previous studies in yeasts have shown that telomeric repeats appended to linear plasmids can promote stable maintenance of linear DNA molecules.^51–53^ We therefore tested whether short terminal repeats derived from *W. ciferrii* could be repurposed to stabilize transient linear donor DNA during homology-independent genome integration.

Mining of *W. ciferrii* sequencing data using Tandem Repeats Finder identified an 11 bp repeat sequence, 5′-CCCAGACACCA-3′. Based on this sequence, we generated VU5 donor variants carrying 0, 1, 2, or 3 tandem copies of the repeat at both termini (**Figure 4A**). Each donor was transformed into the Δ*URA5* recipient strain, and transformants were analyzed using the fluorescence-based VU5 reporter assay. The 2× repeat donor produced the strongest reporter output, with a fluorescence distribution shifted above that of the no-repeat control: 0×, Q1 = 2.4, median = 3.1, Q3 = 3.9; 1×, Q1 = 2.2, median = 2.8, Q3 = 3.9; 2×, Q1 = 3.3, median = 4.9, Q3 = 7.4; and 3×, Q1 = 2.2, median = 2.9, Q3 = 3.5 (**Figure 4B**). The 2× repeat significantly increased fluorescence relative to the 0× donor, whereas the 1× repeat showed no significant improvement and the 3× repeat reduced fluorescence. These results identify two tandem repeats, corresponding to 22 bp per terminus, as the most effective terminal configuration among the variants tested.

**Figure 4.**
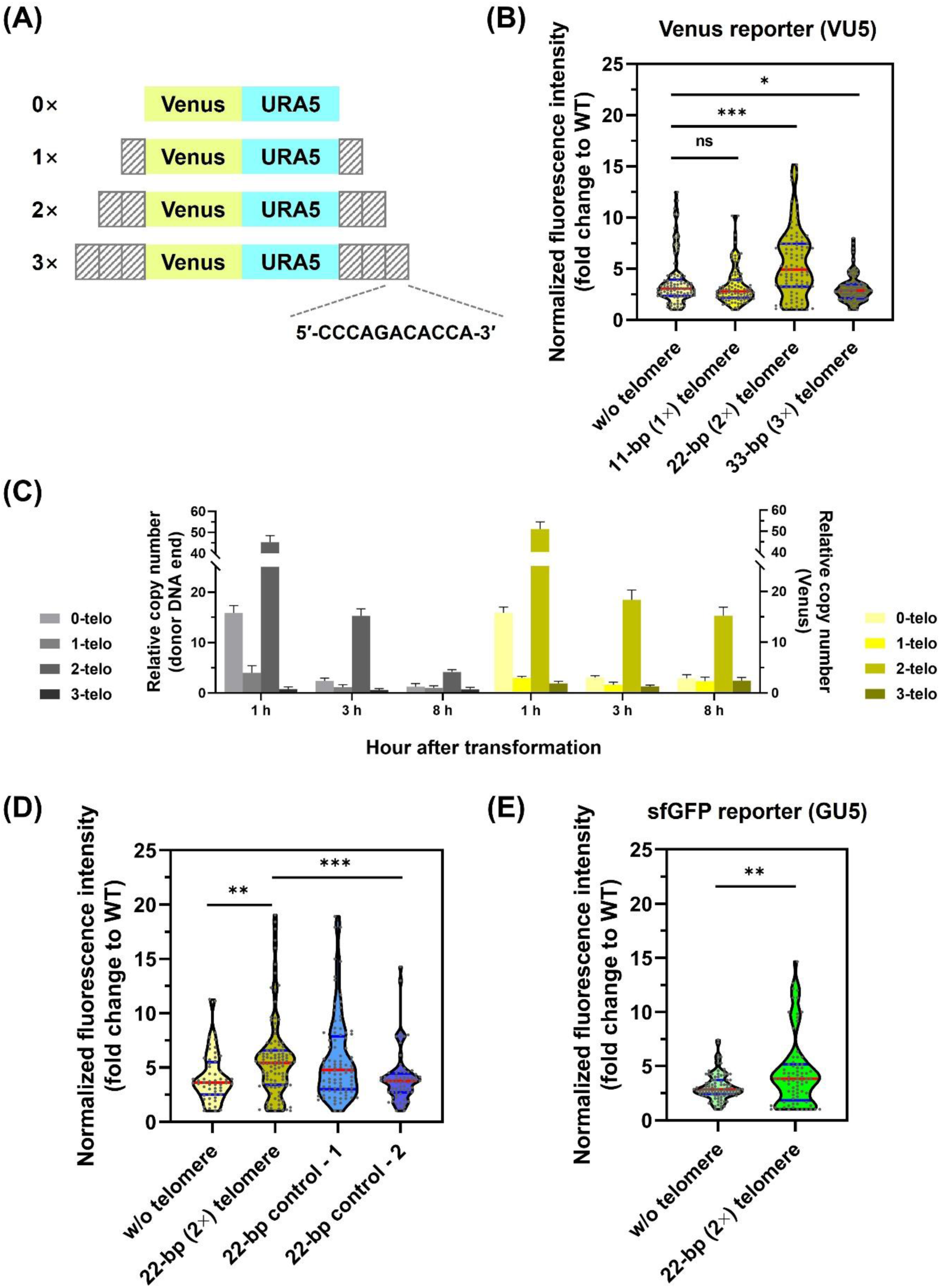
Telomeric end-shielding enhances linear donor stability and random integration output in *W. ciferrii*. **(A)** Design of VU5 donor DNA variants carrying 0, 1, 2, or 3 tandem copies of the 11 bp telomeric repeat sequence, 5′-CCCAGACACCA-3′, at both termini. **(B)** Normalized Venus fluorescence distribution of Δ*URA5* transformants generated with each telomeric donor variant. Fluorescence was normalized to OD_600_ and expressed relative to the wild-type control. **(C)** Time-course qPCR analysis of donor DNA persistence after transformation. Δ*URA5* competent cells were transformed with VU5 donors carrying 0, 1, 2, or 3 telomeric repeats and harvested from the same recovery culture at 1, 3, and 8 h post-transformation. After washing with PBS, donor abundance was quantified using primers targeting either the terminal donor region of the *PGK1* promoter 5′ end or the internal Venus sequence. *TDH3* was used as the diploid genomic reference. Data represent the mean ± SD from three independent transformations. **(D)** Sequence-specific effect of telomeric end-shielding. VU5 donors carrying 2 tandem telomeric repeats were compared with donors carrying two unrelated 22 bp length-matched control sequences at both termini, with the no-telomere donor included as a reference. **(E)** Reporter-independent validation of telomeric end-shielding using a GU5 donor carrying sfGFP instead of Venus. Transformants in **(B)**, **(D)**, and **(E)** were cultured and analyzed as described in Figure 3B. Statistical significance was assessed by Brown-Forsythe and Welch ANOVA followed by Dunnett’s T3 multiple comparisons test relative to the no-telomere control in **(B)** and **(D)**, and by an unpaired *t*-test with Welch’s correction in **(E)**. Red lines indicate medians, and blue lines indicate interquartile ranges.

To determine whether the increased reporter output reflected improved donor DNA persistence after transformation, we quantified donor DNA abundance by qPCR at 1, 3, and 8 h post-transformation (**Figure 4C**). Δ*URA5* competent cells were electroporated with VU5 donors carrying 0, 1, 2, or 3 telomeric repeats, and aliquots were collected from the same recovery culture at each time point. After washing to reduce extracellular DNA carryover, total DNA was extracted and analyzed using primer pairs targeting either the terminal donor region of the *PGK1* promoter or an internal Venus sequence. The terminal signal was corrected for the endogenous *PGK1* a-allele background, and *TDH3* was used as a diploid genomic reference. At 1 h post-transformation, the 2× repeat donor showed the highest apparent abundance for both targets, with approximately 45 copies detected at the donor DNA end and approximately 51 copies detected within Venus. In contrast, the 0× donor showed approximately 16 copies for both regions. At this early time point, qPCR values reflect both cell-associated and integrated donor DNA. By 8 h, the terminal signal declined sharply across all conditions, whereas the internal Venus signal was retained more efficiently in the 2× group, with approximately 15 copies still detectable. These results support a model in which the 2× telomeric repeat improves early donor DNA persistence and helps preserve the internal cassette, although terminal processing still occurs over time.

We next examined whether this effect was specific to the telomeric repeat sequence or could be explained simply by terminal extension length. Two unrelated 22 bp sequences were appended to both termini of the VU5 donor: NC-1, 5′-GGACGCTTCAGCGATACTCCGC-3′, and NC-2, 5′-CCTGCGTCACGGTTGACCGTTC-3′ (**Figure 4D**). The 2× telomeric donor produced significantly higher fluorescence than the no-repeat control and the NC-2 donor. However, NC-1 generated fluorescence levels comparable to those of the 2× telomeric donor. These results indicate that terminal extension itself can contribute to donor protection, but extension length alone does not fully explain the observed effect because the two length-matched control sequences behaved differently. We therefore interpret the 2× telomeric repeat as an empirical optimum that likely combines length-dependent buffering with sequence-dependent properties. Because only two control sequences were tested, the precise sequence features responsible for this difference remain unresolved.

Finally, we tested whether the protective effect of the 2× repeat was dependent on the internal reporter sequence. Venus was replaced with Superfolder green fluorescent protein (sfGFP) to generate the GU5 cassette, and GU5 donors carrying either 0× or 2× terminal repeats were transformed into Δ*URA5* cells. The 2× GU5 donor again showed significantly enhanced fluorescence relative to the 0× donor: 0×, Q1 = 2.4, median = 2.9, Q3 = 3.7; and 2×, Q1 = 2.0, median = 3.8, Q3 = 5.2 (**Figure 4E**). Thus, the benefit of the 2× terminal repeat was reproduced with a distinct internal reporter gene.

Together, these results show that adding two copies of the 11 bp *W. ciferrii* telomeric repeat to each donor terminus improves early linear donor persistence and increases random integration reporter output. Although the effect cannot be attributed exclusively to telomere sequence recognition, the 2× repeat provides a practical and empirically optimized terminal-shielding element for NHEJ-based multicopy genome engineering in *W. ciferrii*.

### Defective *URA5* marker selection enriches multicopy integrants

After improving linear donor performance through telomeric end-shielding, we next sought to increase donor copy number in a single transformation step. NHEJ-mediated random integration can introduce donor DNA without predefined genomic targets or homology arms, but standard auxotrophic markers primarily select for successful complementation and do not necessarily enrich transformants carrying higher copy numbers. We therefore designed a defective-marker strategy in which *URA5* expression was attenuated by promoter truncation. We reasoned that reduced *URA5* transcription would impose stronger selection pressure, thereby favoring transformants that restore uracil prototrophy through integration of multiple donor copies.

Using pVU5 as a template, we generated a series of VU5 donor cassettes in which Venus was linked to *URA5* driven by progressively shortened upstream regions of 500, 100, 20, 10, or 5 bp (**Figure 5A**). Each cassette was cloned into a plasmid, PCR-amplified as a linear donor fragment, and transformed into Δ*URA5* cells. Colony recovery decreased markedly as the *URA5* promoter was shortened, consistent with increased selection stringency. The 5 bp promoter variant yielded too few transformants for reliable fluorescence-based screening and was therefore excluded from quantitative analysis.

**Figure 5.**
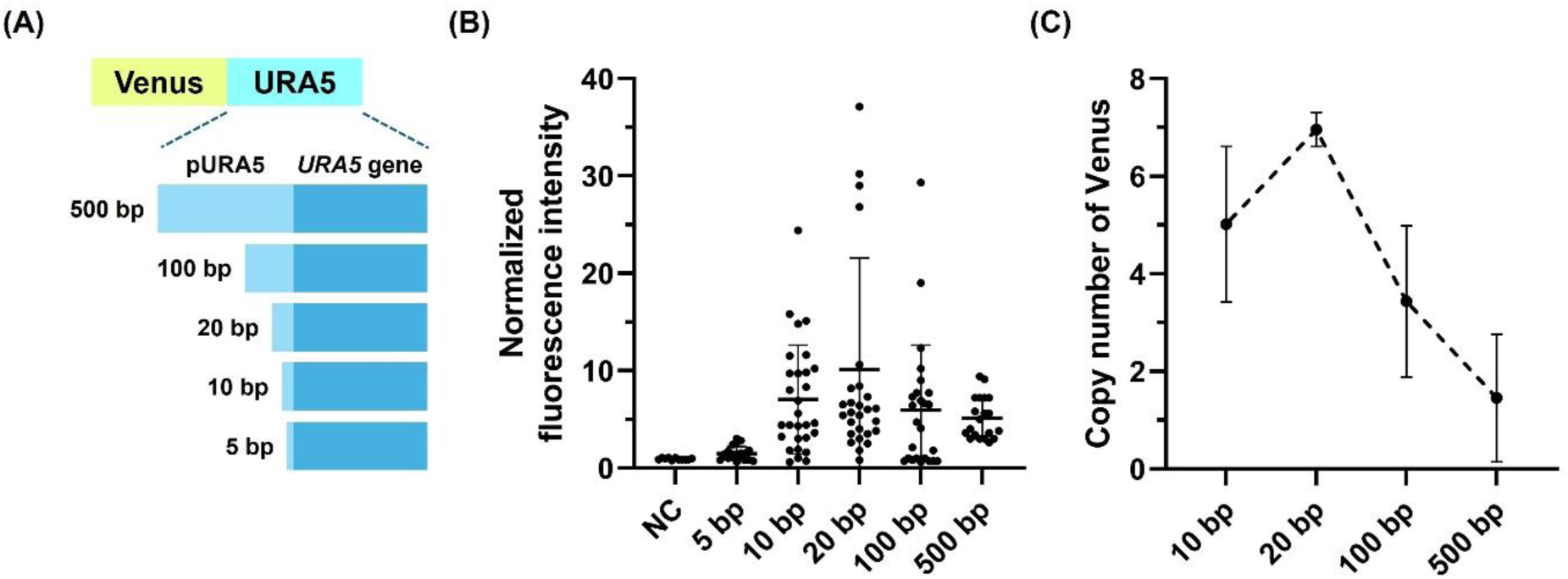
Defective *URA5* marker selection enriches multicopy random integrants in *W. ciferrii*. **(A)** Design of VU5 donor variants carrying Venus linked to *URA5* expression cassettes with progressively truncated *URA5* promoter regions of 500, 100, 20, 10, or 5 bp. **(B)** Normalized Venus fluorescence of transformants recovered from each promoter-length condition. Twenty-five integrants per condition were cultivated in YMglSC medium in 96-deep-well plates for 48 h before fluorescence measurement. NC indicates the non-fluorescent Δ*URA5* negative control. Horizontal bars indicate the mean. **(C)** Venus copy number in the three highest-fluorescence clones from each promoter group, determined by qPCR using *TDH3* as a diploid genomic reference. Data represent the mean ± SD from three selected clones.

To compare reporter output among the promoter variants, 25 integrants from each recoverable condition were cultivated in 96-deep-well plates containing YMglSC medium, and Venus fluorescence was measured after 48 h. Fluorescence was normalized to OD_600_ and expressed relative to the non-fluorescent control. Among the tested constructs, the 20 bp *URA5* promoter produced the highest fluorescence distribution (**Figure 5B**), suggesting that intermediate marker attenuation most effectively enriched transformants with elevated donor dosage. To validate this interpretation, we quantified donor copy number by qPCR using the three highest-fluorescence clones from each promoter group. The estimated donor copy number increased from 1.5 copies for the 500 bp promoter to 3.4 copies for the 100 bp promoter and 7.0 copies for the 20 bp promoter, followed by a decrease to 5.0 copies for the 10 bp promoter (**Figure 5C**). Because qPCR was performed on the highest-fluorescence clones, these values represent enriched integrants rather than population-wide copy-number averages.

These results show that attenuating *URA5* expression converts uracil prototrophy into a copy-number-responsive selection pressure in *W. ciferrii*. The 20 bp *URA5* promoter provided the most favorable balance between transformant recovery and multicopy enrichment, whereas further truncation appeared to impose excessive stringency that limited productive recovery. Thus, defective *URA5* marker selection provides a practical strategy for enriching multicopy NHEJ-mediated integrants in a single transformation step.

### Donor-end configuration enhances transformant recovery and random integration output

Having established telomeric end-shielding and defective *URA5* selection as two design principles for improving random integration, we next examined whether donor-end configuration further influenced NHEJ-mediated genome insertion. Because NHEJ requires ligation-compatible DNA termini, we reasoned that both the chemical state and structure of donor ends could affect transformant recovery and integration output.

To test this, we generated three types of linear VU5 donor fragments with distinct end configurations (**Figure 6A**). The first was an unmodified PCR product amplified from pVU5 using primers pPGK1_F and Gibson_DM_tURA5_R, which generated non-phosphorylated 5′ termini. The second was a sticky-end donor amplified with primers XbaI_pPGK1_F and XSmaI_tURA5_R and then digested with XbaI and XmaI to generate restriction enzyme-processed cohesive termini with 5′ phosphorylation. The third was a blunt-end donor amplified with primers SmaI_pPGK1_F and XSmaI_tURA5_R and digested with SmaI to generate restriction enzyme-processed blunt termini with 5′ phosphorylation (**Table 2**). Each donor was transformed into Δ*URA5* cells, and transformant recovery was quantified on SD-Ura agar.

**Figure 6.**
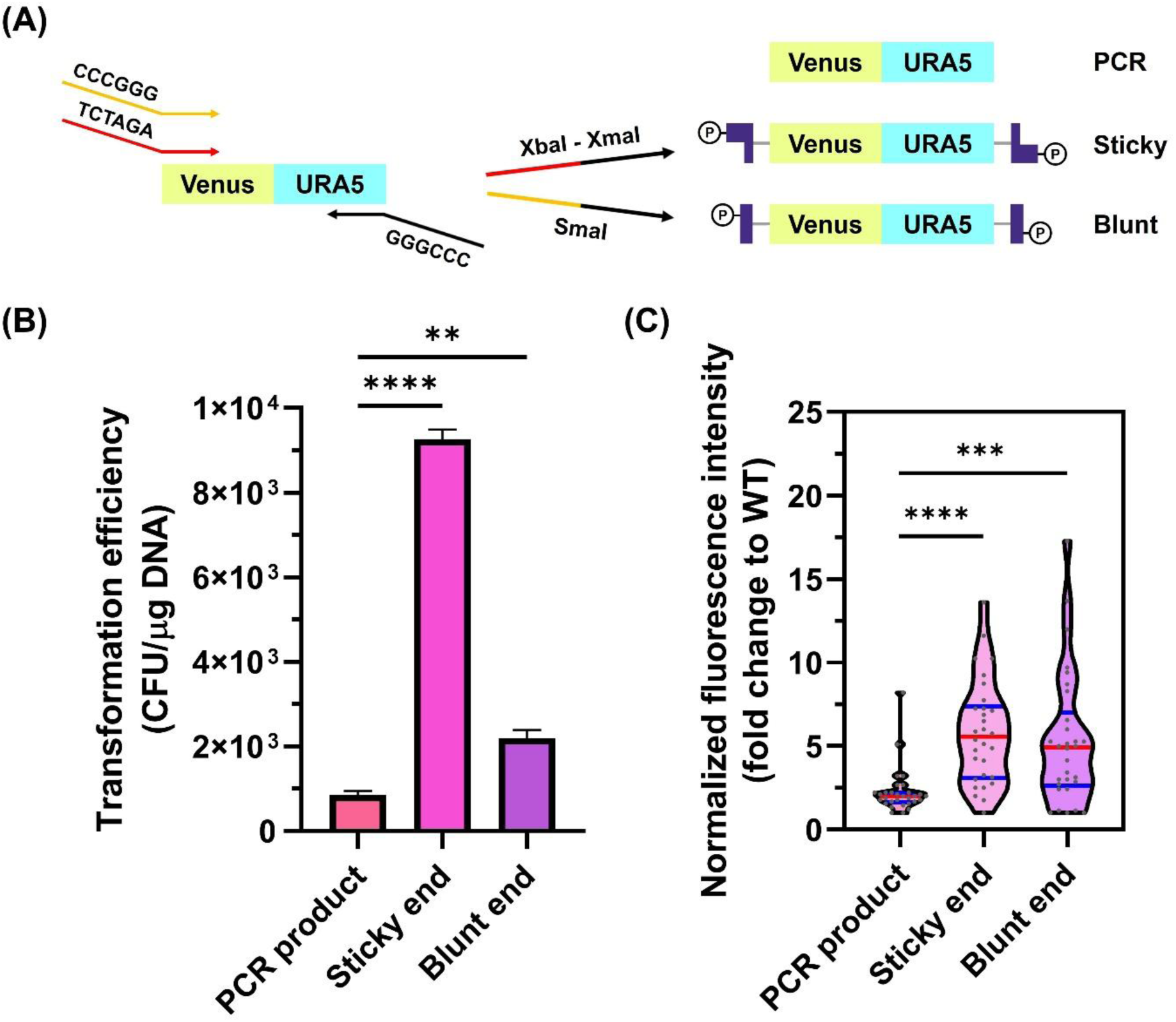
Donor-end configuration influences transformant recovery and random integration output in *W. ciferrii*. **(A)** Schematic of the three VU5 donor DNA types used to evaluate end-configuration effects: an unmodified PCR product with non-phosphorylated termini, a sticky-end donor generated by XbaI–XmaI digestion, and a blunt-end donor generated by SmaI digestion. Restriction enzyme-generated donors carry 5′-phosphorylated termini (circled P). **(B)** Transformation efficiency of each donor type, expressed as colony-forming units (CFU) per μg DNA on SD-Ura agar. Data represent the mean ± SD from three independent transformations. **(C)** Normalized Venus fluorescence distribution of transformants obtained with each donor type. Thirty-two transformants per condition were cultivated and analyzed as described in Figure 3B. Statistical significance was assessed by Brown-Forsythe and Welch ANOVA followed by Dunnett’s T3 multiple comparisons relative to the unmodified PCR product. Red lines indicate medians, and blue lines indicate interquartile ranges.

Donor-end configuration had a pronounced effect on transformation efficiency. Non-phosphorylated PCR products yielded 821 ± 86 CFU/μg DNA, whereas sticky-end donors yielded 9,168 ± 226 CFU/μg DNA and blunt-end donors yielded 2,118 ± 190 CFU/μg DNA (**Figure 6B**). Thus, restriction enzyme-generated donors substantially improved transformant recovery relative to unmodified PCR products, with approximately 11-fold higher recovery for sticky-end donors and 2.6-fold higher recovery for blunt-end donors. The higher recovery of sticky-end fragments compared with blunt-end fragments further suggests that donor-end structure influences early transformation or integration outcomes.

We next assessed whether donor-end configuration also affected reporter output among recovered integrants using the VU5 fluorescence assay. For each donor type, 32 transformants and three non-fluorescent controls were cultivated in 96-deep-well plates containing YMglSC medium for 48 h. Fluorescence was normalized to OD_600_ and expressed relative to the non-fluorescent control. After outlier removal, the fluorescence distributions were as follows: PCR products, Q1 = 1.7, median = 2.0, Q3 = 2.2; sticky end, Q1 = 3.1, median = 5.6, Q3 = 7.3; and blunt end, Q1 = 2.7, median = 4.9, Q3 = 6.6 (**Figure 6C**). Both sticky-end and blunt-end donors produced significantly higher fluorescence than the PCR product, whereas no significant difference was observed between the sticky-end and blunt-end groups. These results indicate that restriction enzyme-generated, 5′-phosphorylated termini improve reporter output among recovered integrants, while cohesive and blunt termini perform similarly once stable transformants are obtained.

Mechanistically, this effect is consistent with the requirements of canonical NHEJ ligation. The DNA ligase IV complex catalyzes phosphodiester bond formation between a 5′-phosphate and a 3′-hydroxyl group at DNA break site.^54^ Accordingly, non-phosphorylated PCR donor ends likely require additional processing before ligation, whereas restriction enzyme-generated donors already carry ligation-compatible 5′-phosphorylated termini.^55,56^ Although sticky ends improved overall transformant recovery more strongly than blunt ends, the comparable fluorescence distributions of sticky- and blunt-end integrants suggest that 5′-phosphorylated, enzyme-processed termini are sufficient to enhance random integration output in *W. ciferrii*. Together, these findings identify donor-end optimization as a third design parameter for improving NHEJ-based genome integration and support the use of restriction enzyme-released, 5′-phosphorylated donor fragments in the final platform vector.

### Multicopy integration of *LCB1* and *LCB2* enhances TAPS production

Having optimized telomeric end-shielding, defective *URA5* marker selection, and 5′-phosphorylated donor termini, we consolidated these design elements into a single platform vector for pathway engineering in *W. ciferrii*. The resulting vector, pTdmVU5, contains terminal multiple cloning sites for donor release, two tandem copies of the 11 bp *W. ciferrii* telomeric repeat at each terminus, MoClo-compatible connector sequences, a SwaI site for Gibson assembly-based cassette insertion, and a loxP-flanked defective VU5 selection module composed of Venus and *URA5* driven by the 20 bp truncated *URA5* promoter (**Figure 7A**). This architecture was designed to enable modular cassette assembly, release of 5′-phosphorylated linear donor DNA, fluorescence-based screening, and potential marker recycling.

**Figure 7.**
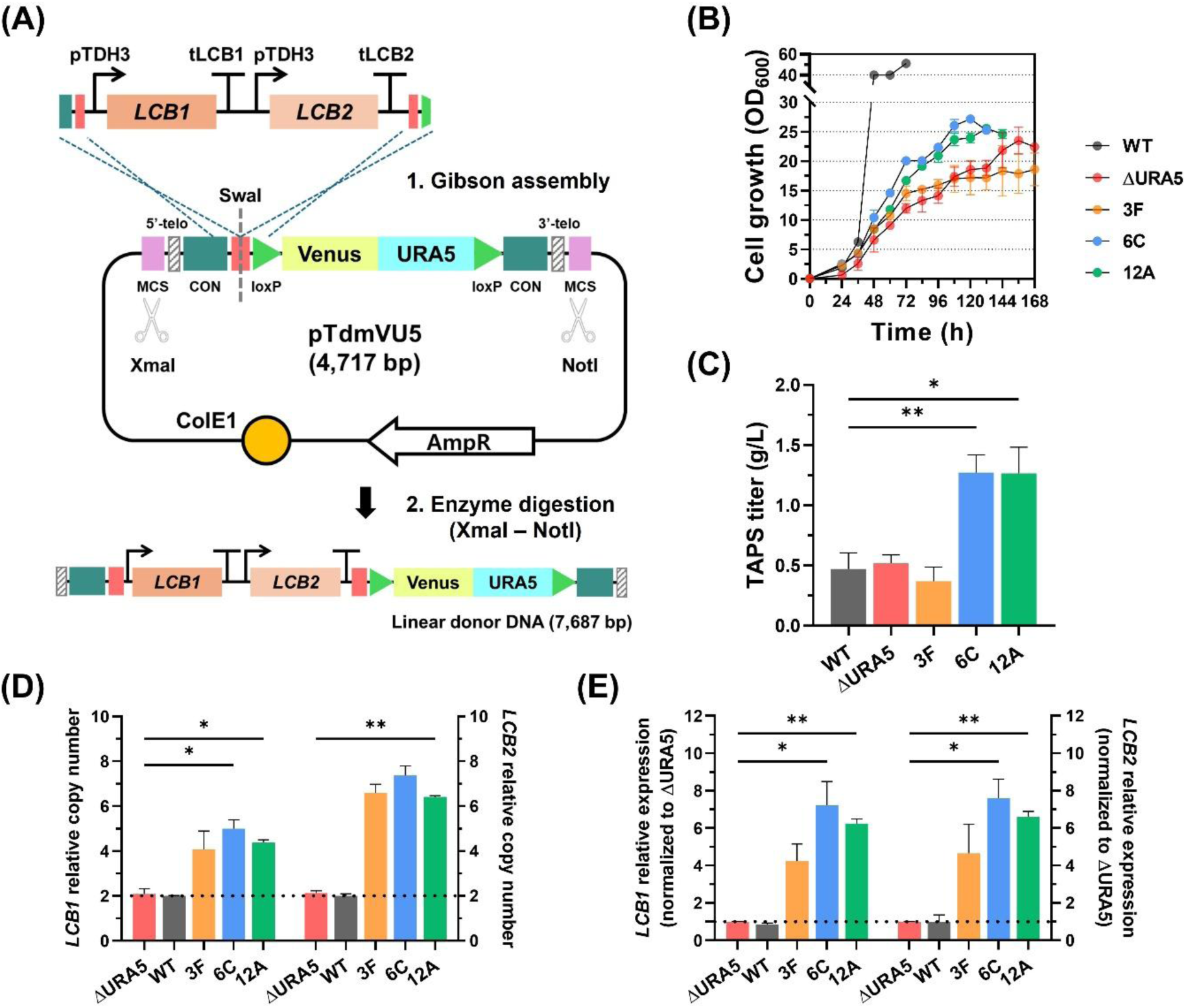
Multicopy integration of *LCB1* and *LCB2* using pTdmVU5 enhances TAPS production in *W. ciferrii*. **(A)** Schematic of pTdmVU5-mediated donor construction. Two expression cassettes, pTDH3-*LCB1*-tLCB1 and pTDH3-*LCB2*-tLCB2, were inserted into the SwaI site by Gibson assembly. The complete 7,687 bp donor fragment was released by XmaI–NotI digestion for transformation. Key platform elements include terminal telomeric repeats, MoClo connector sequences (CON), loxP sites flanking the defective VU5 module, and multiple cloning sites (MCS). **(B)** Growth profiles of wild-type *W. ciferrii* (WT), the parental Δ*URA5* strain, and three recovered integrant strains, 3F, 6C, and 12A, cultivated in AAT medium at 25 °C. **(C)** TAPS titers at the end of cultivation. **(D)** Genomic copy numbers of *LCB1* and *LCB2* determined by qPCR using *TDH3* as a diploid genomic reference. Values include endogenous and integrated copies; the dotted line indicates the endogenous diploid copy number. **(E)** Relative transcript levels of *LCB1* and *LCB2* determined by RT-qPCR using *ACT1* as the reference gene and the ΔΔCt method. Expression levels were normalized to the parental Δ*URA5* strain; the dotted line indicates the basal expression level in Δ*URA5*. Data represent the mean ± SD from three biological replicates. Statistical significance was assessed by Brown-Forsythe and Welch ANOVA followed by Dunnett’s T3 multiple comparisons test, relative to the wild-type control in **(C)** and the parental Δ*URA5* control in **(D)** and **(E)**.

To evaluate the platform in a metabolic engineering context, we targeted the first committed step of sphingolipid biosynthesis. *LCB1* and *LCB2*, which encode the two subunits of serine palmitoyltransferase, were selected because this enzyme complex controls sphingoid base flux in *W. ciferrii* (**Figure S1**).^24^ Two expression cassettes, pTDH3-*LCB1*-tLCB1 and pTDH3-*LCB2*-tLCB2, were assembled into SwaI-linearized pTdmVU5 by Gibson assembly, generating pTdmVU5-pTDH3_*LCB1*-pTDH3_*LCB2* (**Figure 7A**). The complete donor fragment was released by XmaI–NotI digestion as an approximately 7.7 kb sticky-end linear cassette and transformed into Δ*URA5* cells.

Direct selection on SD-Ura agar yielded very few colonies, in contrast to the shorter VU5 reporter donor. This result indicates that donor length imposes a substantial constraint on NHEJ-mediated random integration. The stringent 20 bp defective *URA5* promoter likely further amplified this limitation because colony formation requires sufficient integrated copy number to restore uracil prototrophy. Thus, long donor molecules that integrate at sub-threshold copy numbers may not be recovered under direct auxotrophic selection.

To overcome this recovery bottleneck, we used Venus fluorescence as a selection-independent signal for isolating integrants. Following transformation, cells were cultured overnight in YPD medium to allow outgrowth, and Venus-positive cells were isolated by fluorescence-activated cell sorting (FACS) and plated on YPD agar. This strategy enabled recovery of integrants regardless of uracil prototrophy. Ninety-six colonies were then screened in 96-deep-well plates, and the three clones with the highest normalized fluorescence, designated 3F, 6C, and 12A, were selected for further characterization.

The selected strains, together with wild-type *W. ciferrii* and the parental Δ*URA5* strain, were cultivated in baffled flasks containing AAT medium. Growth kinetics differed markedly among the strains (**Figure 7B**). The wild-type strain consumed 30 g/L glycerol within 70 h and reached an OD_600_ of approximately 40. In contrast, Δ*URA5* and strain 3F required 168 h to exhaust glycerol and reached final OD_600_ of approximately 22 and 19, respectively. Strains 6C and 12A showed intermediate growth kinetics, consuming glycerol within 120–130 h and reaching OD_600_ values of approximately 25.

TAPS production confirmed the metabolic utility of the integration platform (**Figure 7C**). Wild-type and Δ*URA5* strains produced 0.47 and 0.52 g/L TAPS, respectively, indicating that loss of *URA5* did not substantially reduce final TAPS titer under these cultivation conditions, although it slowed growth and glycerol utilization. Strain 3F produced 0.37 g/L TAPS, the lowest titer among the tested strains, suggesting that its random integration event may have imposed a fitness or pathway-specific burden. In contrast, strains 6C and 12A produced 1.27 and 1.26 g/L TAPS, respectively, corresponding to approximately 2.7-fold higher titers than the wild-type strain. These results demonstrate that pTdmVU5-mediated integration can rapidly generate productive multicopy pathway-amplified strains.

We next quantified integrated donor copy number by qPCR using *TDH3* as a diploid genomic reference (**Figure 7D**). Wild-type and Δ*URA5* strains contained approximately two copies each of endogenous *LCB1* and *LCB2*, as expected for diploid cells. In the integrant strains, estimated copy number increased to 4.1 and 6.6 copies of *LCB1* and *LCB2* in strain 3F, 5.0 and 7.4 copies in strain 6C, and 4.4 and 6.4 copies in strain 12A, respectively. Across all integrants, *LCB2* copy number was higher than *LCB1* copy number. Because *LCB1* is positioned closer to one donor terminus whereas *LCB2* is located more internally, this imbalance is consistent with partial terminal processing of long linear donor molecules before or during integration. Thus, although telomeric end-shielding improves donor persistence, it does not completely eliminate end resection.

RT-qPCR confirmed that increased gene dosage was accompanied by elevated transcript abundance (**Figure 7E**). Wild-type and Δ*URA5* strains showed no significant difference in *LCB1* or *LCB2* transcript levels, indicating that *URA5* disruption did not itself alter expression of these sphingolipid biosynthetic genes. Relative to the parental Δ*URA5* strain, strain 3F showed 4.2- and 4.7-fold increases in *LCB1* and *LCB2* transcript levels, respectively; strain 6C showed 7.2- and 7.7-fold increases; and strain 12A showed 6.2- and 6.6-fold increases. Transcript abundance therefore broadly paralleled donor copy number. The high transcript levels and increased TAPS titers in strains 6C and 12A support the conclusion that multicopy integration of *LCB1* and *LCB2* enhances sphingoid base flux. However, the similar titers of 6C and 12A despite different copy numbers and transcript levels suggest that TAPS production begins to encounter a downstream bottleneck at these expression levels. Further improvement will likely require combinatorial engineering of additional nodes, such as competing phosphorylation, hydroxylation, acetylation, or precursor supply.

The long *LCB1*–*LCB2* donor also revealed a practical limitation of the defective-marker system. Although the integrant strains carried approximately 4–5 copies of the terminally positioned *LCB1* cassette, none recovered growth on SD-Ura agar. In the shorter VU5 donor experiments, recovery on SD-Ura agar was associated with approximately seven copies for the 20 bp *URA5* promoter, suggesting an approximate copy-number threshold for sufficient marker expression. The longer donor therefore appears to integrate below the threshold required for direct defective-marker selection, explaining the poor colony recovery on SD-Ura agar. Nevertheless, strains 6C and 12A showed partially improved growth compared with the Δ*URA5* parent under AAT cultivation, suggesting that multiple defective *URA5* copies provide low but detectable orotate phosphoribosyltransferase activity under non-dropout medium conditions.

Together, these results demonstrate that the pTdmVU5 platform can generate multicopy pathway-amplified *W. ciferrii* strains and increase TAPS production through simultaneous integration of *LCB1* and *LCB2*. At the same time, the long-donor experiment revealed a practical limitation: the stringent 20 bp defective *URA5* promoter, although effective for multicopy enrichment of compact cassettes, becomes restrictive for larger pathway constructs that integrate at sub-threshold copy numbers. To accommodate such donors, we also constructed a companion vector, pToriVU5, in which the 20 bp truncated *URA5* promoter was replaced with the full-length 500 bp *URA5* promoter; preliminary tests using a heterologous donor cassette of approximately 8 kb showed robust colony recovery on SD-Ura agar, suggesting that pTdmVU5 and pToriVU5 may serve as complementary tools for compact and larger donor-size regimes, respectively.

## Materials and methods

### Strains, media, and growth conditions

*Wickerhamomyces ciferrii* (F-60-10A NRRL 1031) was used as the parental strain throughout this study. *Saccharomyces cerevisiae* (CEN.PK2-1D) was used as a control for the split-fluorescent reporter assay. *Escherichia coli* NEB 10-beta (New England Biolabs, Ipswich, MA, USA) served as the host for plasmid construction and propagation. All strains and plasmids used in this study are listed in **Table 1**.

**Table 1.**
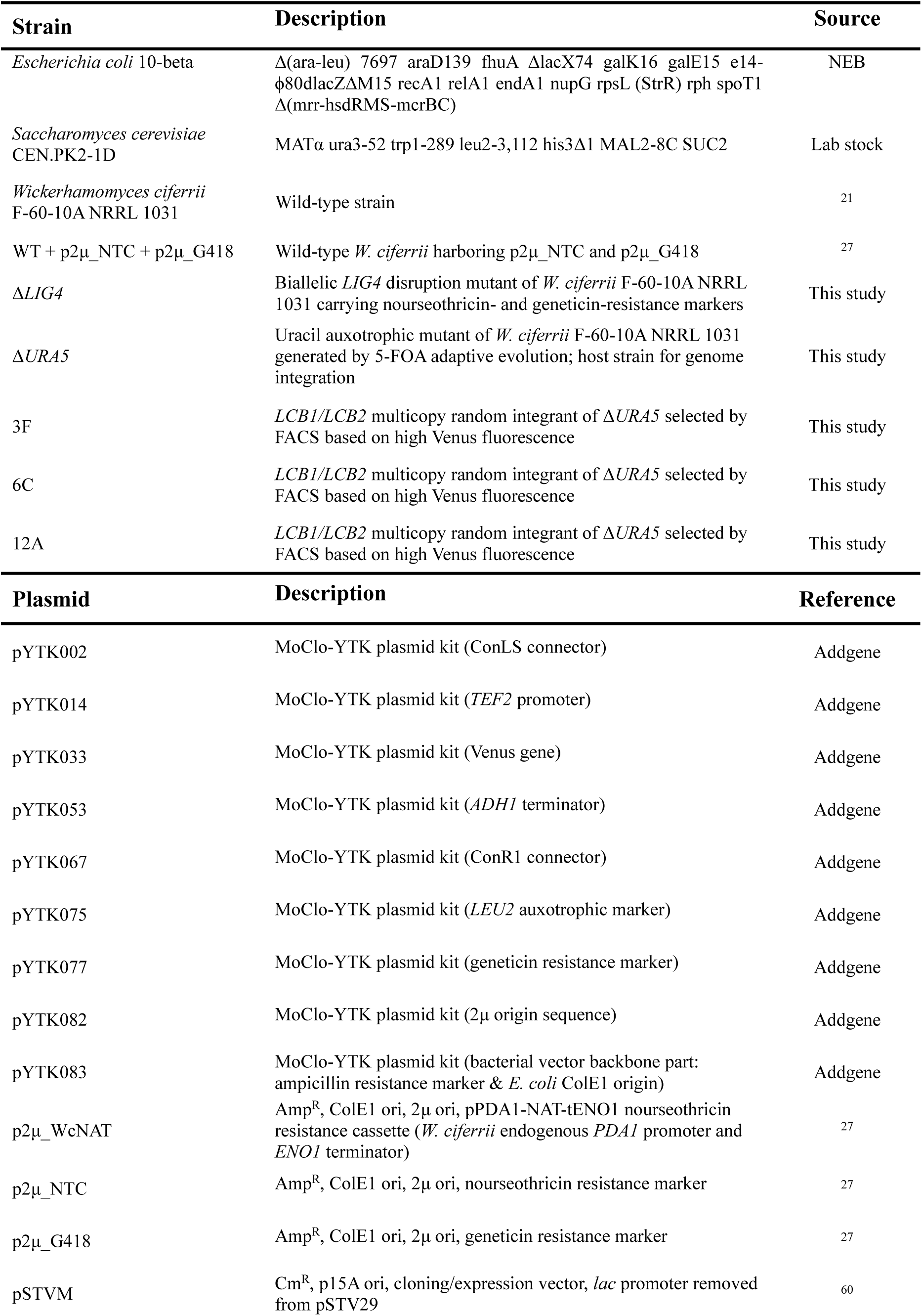
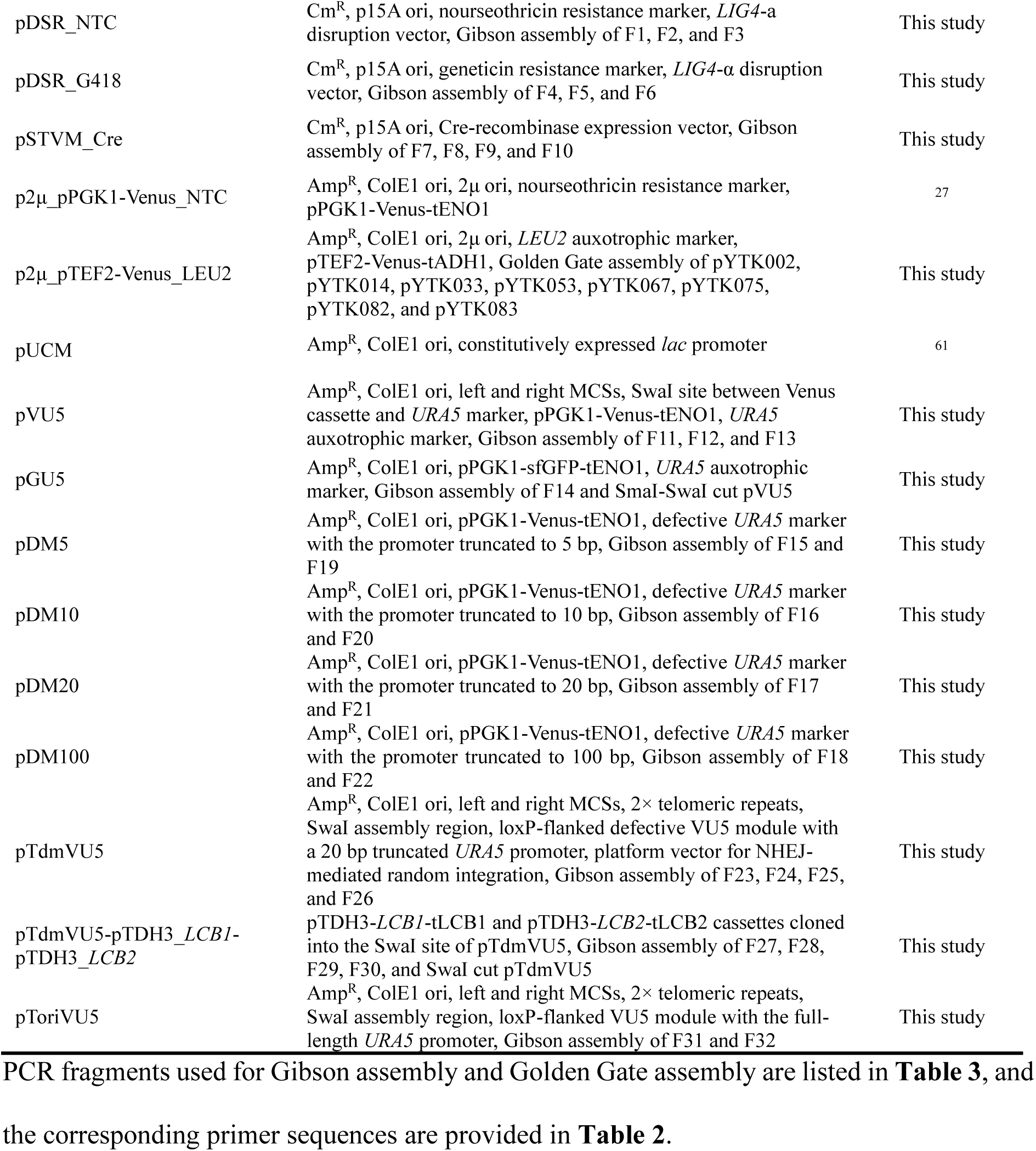
Strains and plasmids used in this study.

*W. ciferrii* was routinely grown on YPD agar (10 g/L yeast extract, 20 g/L peptone, 20 g/L glucose, and 15 g/L agar) at 25 °C. For selection of antibiotic-resistant transformants, YPD agar was supplemented with 100 μg/mL nourseothricin or 300 μg/mL geneticin. Liquid cultivations were performed in the following media depending on experimental purpose: YPD medium (10 g/L yeast extract, 20 g/L peptone, and 20 g/L glucose) for routine growth and competent cell preparation; YMglSC medium (2 g/L yeast extract, 2 g/L malt extract, 7 g/L peptone, 20 g/L glycerol, 10 mM CaCl_2_, and 5 g/L serine) for short-term cultivation including fluorescence screening, copy number determination, and transcriptional analysis; and AAT medium (3 g/L yeast extract, 1 g/L KNO_3_, 1 g/L KH_2_PO_4_, 4 g/L sodium glutamate, 1 g/L glycine, 3 g/L sodium acetate, 0.5 g/L MgSO_4_·7H_2_O, 1 g/L CaCl_2_·2H_2_O, 25 mg/L uracil, and 30 g/L glycerol) for flask-scale TAPS production. For liquid cultivation of antibiotic-resistant strains, 50 μg/mL nourseothricin or 50 μg/mL geneticin was added. SD-Ura agar (synthetic defined medium lacking uracil) was used for selection of uracil prototrophs (6.7 g/L yeast nitrogen base (Difco, Sparks, MD, USA), a mixture of amino acids and nucleotides except uracil (synthetic drop-out medium supplements, Sigma-Aldrich, St. Louis, MO, USA), 20 g/L glucose, and 15 g/L agar). 5-Fluoroorotic acid (5-FOA) agar (7 g/L yeast nitrogen base, 1 g/L 5-FOA, 50 mg/L uracil, 20 g/L glucose, and 15 g/L agar) was used for counterselection during uracil auxotroph construction and screening. Unless otherwise stated, *W. ciferrii* was grown at 25 °C with shaking at 250 rpm for shake-flask cultivation or 800 rpm for 96-deep-well plate cultivation.

*E. coli* was grown aerobically at 37 °C with shaking at 250 rpm in LB medium (10 g/L tryptone, 5 g/L yeast extract, and 5 g/L NaCl). 100 μg/mL ampicillin was added for plasmid selection. Cell growth was monitored by measuring the optical density at 600 nm (OD_600_) using a SPECTRAmax PLUS384 spectrophotometer (Molecular Devices, San Jose, CA, USA).

### Genomic DNA extraction

Genomic DNA was extracted from *W. ciferrii* following the protocol of Dymond et al.^57^ with minor modifications, as described previously.^27^ Briefly, a single colony from YPD agar was inoculated into 2 mL of YPD medium and cultured overnight at 25 °C with shaking at 250 rpm. Cells from 1 mL of culture were harvested by centrifugation at 14,000 rpm for 1 min at room temperature and resuspended in 200 μL of yeast lysis buffer containing 10 mM Tris-HCl (pH 8.0), 1 mM EDTA (pH 8.0), 100 mM NaCl, 2% Triton X-100, and 1% SDS. 150 μL of glass beads and 200 μL of phenol/chloroform/isoamyl alcohol (25:24:1, v/v/v) were added, and the mixture was vortexed vigorously for 10 min. 400 μL of TE buffer containing 10 mM Tris-HCl (pH 8.0) and 1 mM EDTA (pH 8.0) was then added, followed by an additional 1 min of vortexing. The mixture was centrifuged at 13,000 rpm for 10 min at 4 °C, and the aqueous phase was transferred to a new tube. DNA was precipitated by adding an equal volume of isopropyl alcohol and mixing by inversion, followed by centrifugation at 13,000 rpm for 10 min at 4 °C. The pellet was washed with 1 mL of ice-cold 70% ethanol, air-dried at room temperature for approximately 5 min after removal of the supernatant, and resuspended in 100 μL of TE buffer.

### Construction of the Δ*URA5* recipient strain

A uracil auxotrophic mutant of the diploid *W. ciferrii* was generated by adaptive evolution on 5-FOA as previously described.^25^ Wild-type cells were serially cultured in YPD medium containing 50 mg/L uracil and stepwise increasing concentrations of 5-FOA from 0.1 to 2.5 g/L. At each concentration step, cultures showing visible growth were streaked onto 5-FOA agar to isolate auxotrophic mutants. Resulting isolates were re-streaked onto SD-Ura agar to screen for uracil auxotrophy. Colonies that failed to grow on SD-Ura agar were selected as candidate auxotrophs, whereas wild-type controls grew normally.

Disruption or loss of the *URA5* locus was examined by genomic PCR. Using primers pURA5_F and Gibson_DM_tURA5_R (**Table 2**), the wild-type strain produced the expected amplicon corresponding to the intact *URA5* locus, whereas no amplicon was detected in the selected candidates. For complementation, a *URA5* cassette containing 500 bp upstream and 140 bp downstream of the coding sequence was PCR-amplified using the same primer pair and transformed into the candidate strain. Successful complementation restored growth on SD-Ura agar and abolished 5-FOA resistance, confirming that the auxotrophic phenotype resulted from loss of *URA5* function. The resulting Δ*URA5* strain was used as the recipient strain for subsequent genome integration experiments.

**Table 2.**
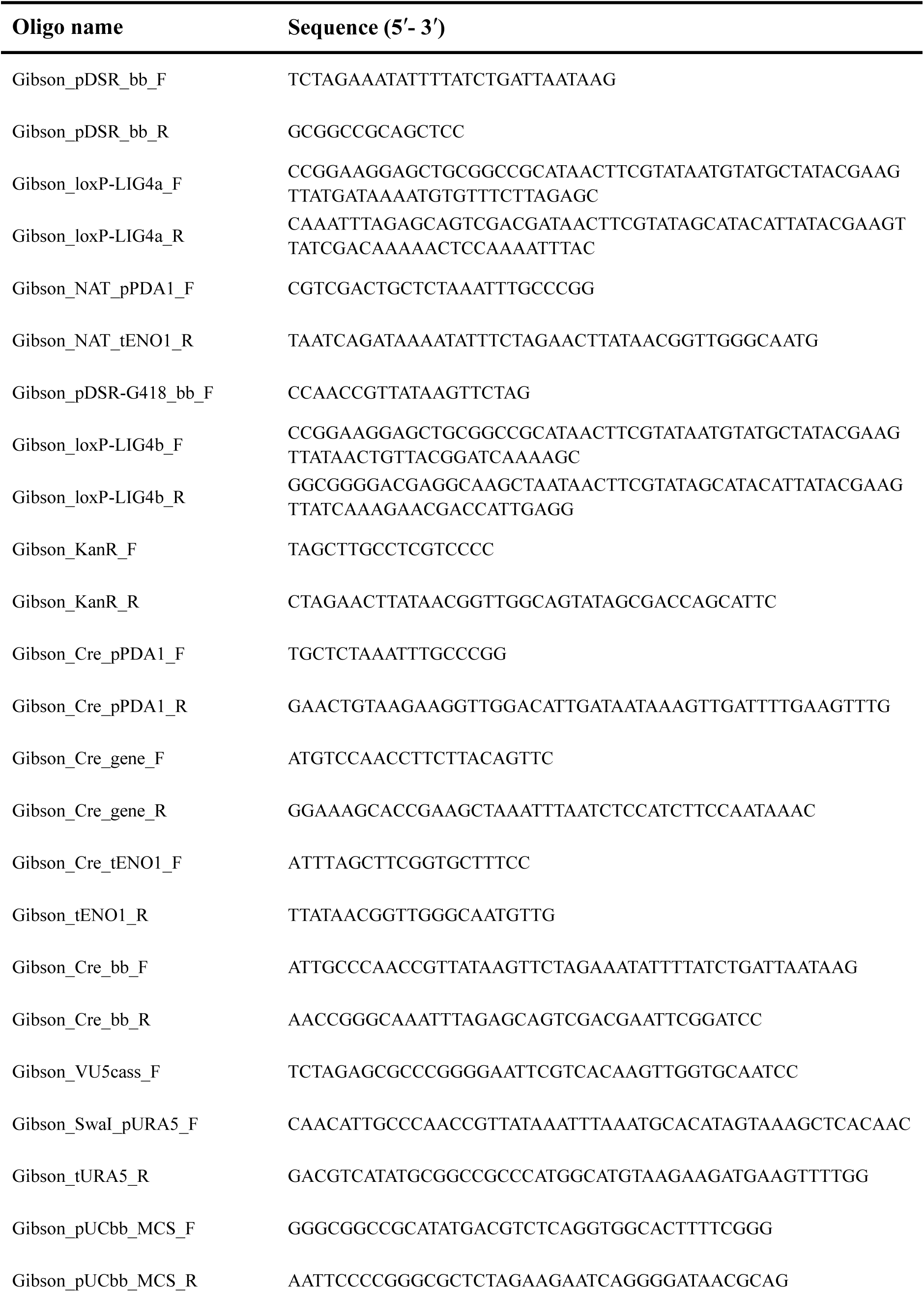
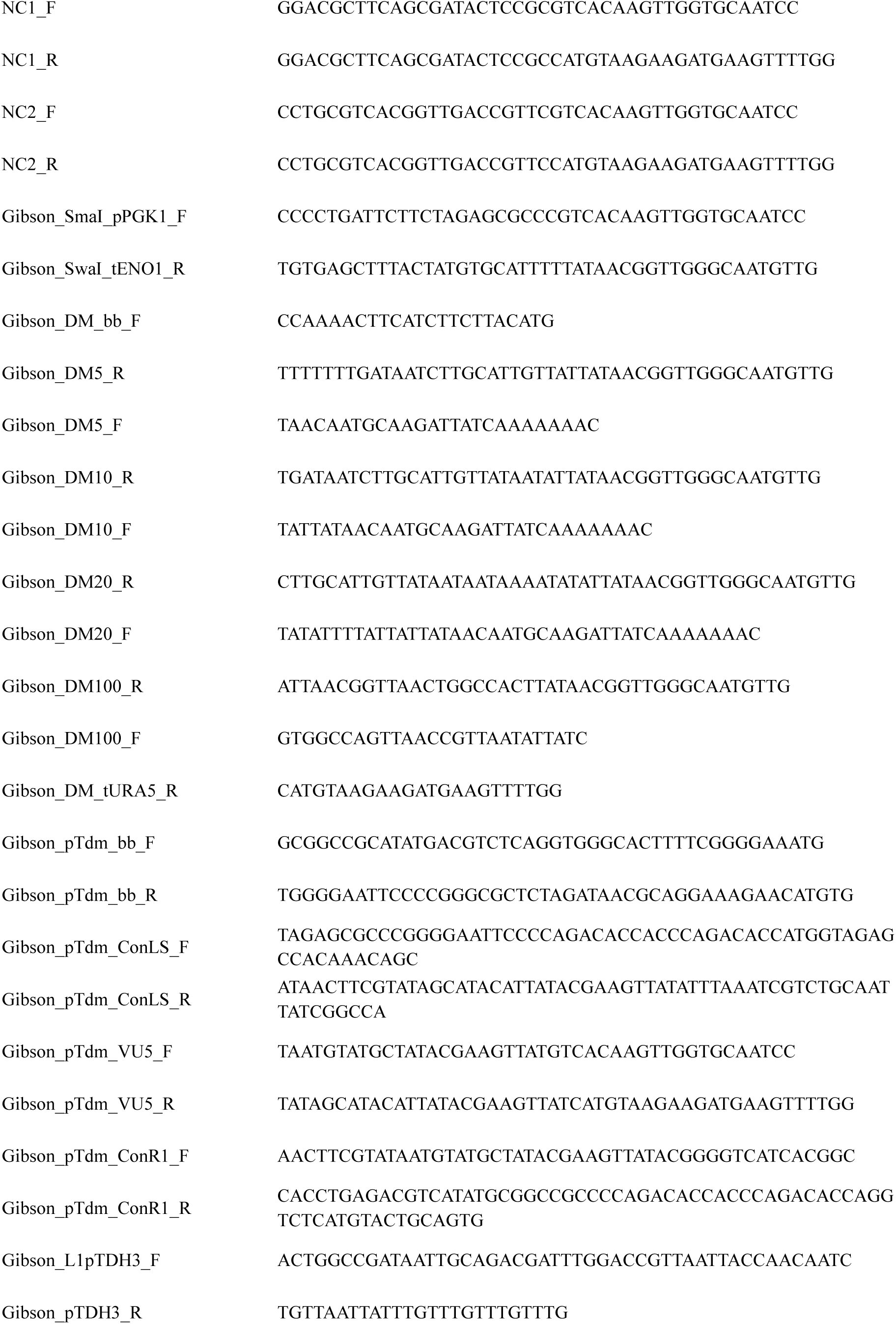
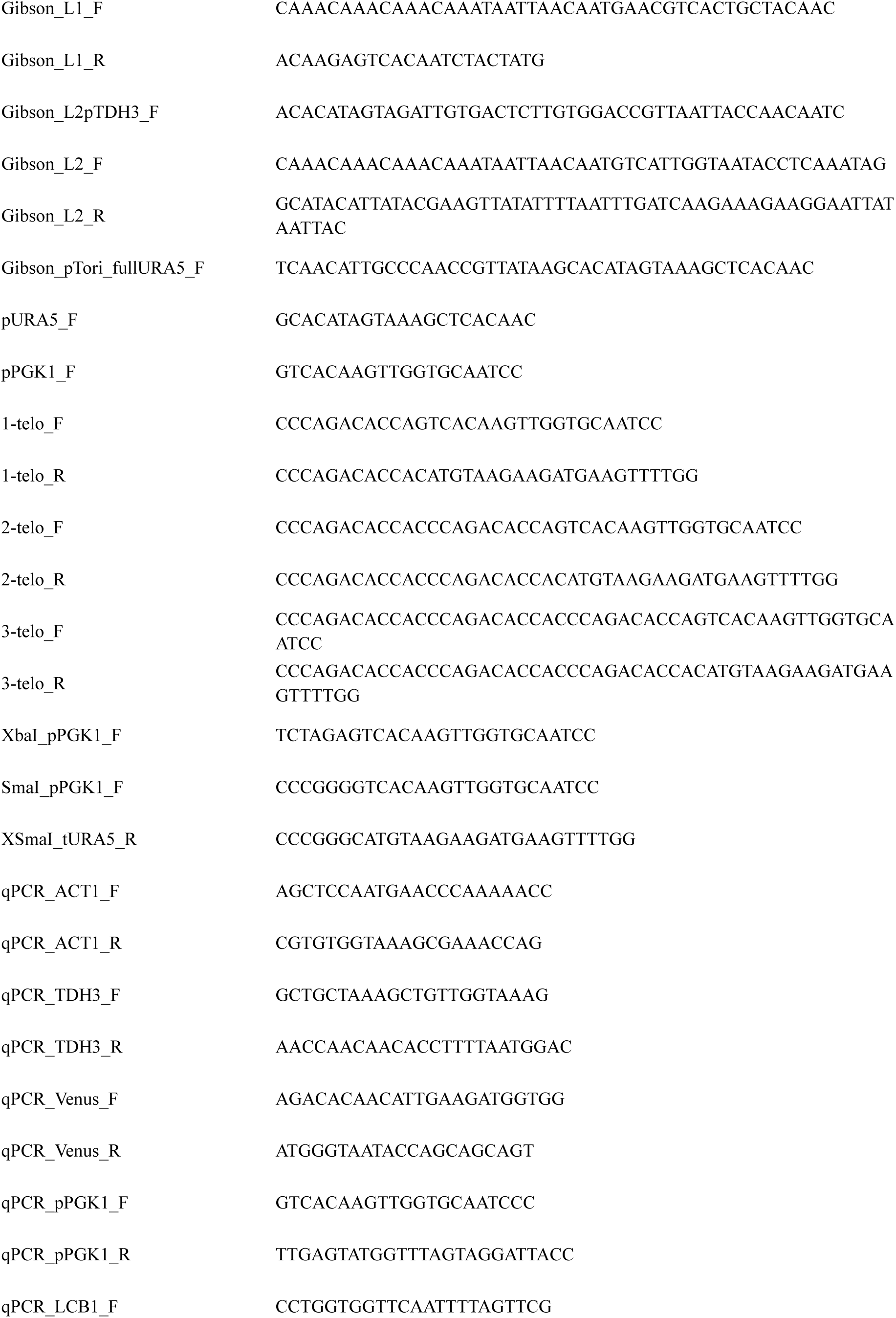
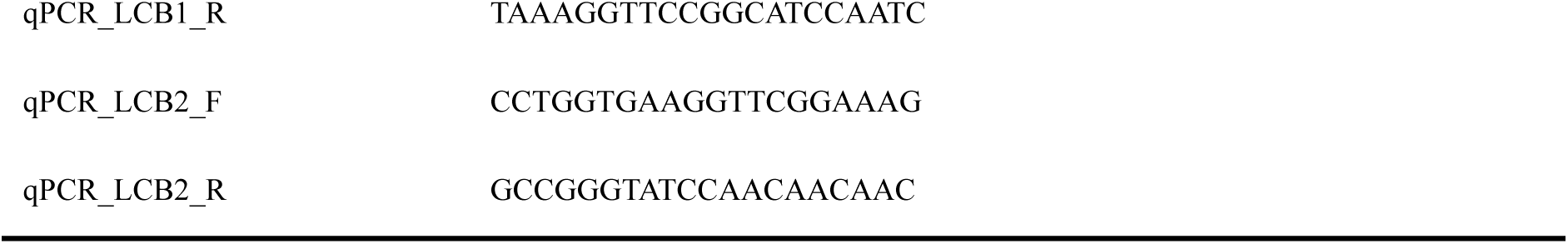
Primers used in this study.

### General DNA assembly and plasmid construction

All plasmids and primers used in this study are listed in **Tables 1** and **2**, respectively. DNA fragments were amplified using Phusion High-Fidelity DNA polymerase (New England Biolabs, Ipswich, MA, USA) and purified using a gel extraction kit (GeneAll Biotechnology, Seoul, Republic of Korea). Oligonucleotides were synthesized by Macrogen (Seoul, Republic of Korea). Plasmids were constructed by Gibson assembly^58^ or Golden Gate assembly.

Gibson assembly was performed using NEBuilder® HiFi DNA Assembly Master Mix (New England Biolabs, Ipswich, MA, USA), and molar ratios were calculated using the NEBuilder Protocol Calculator (https://nebuildercalculator.neb.com/). For Golden Gate assembly, DNA parts were mixed equimolarly at 20 fmol each with 20 U of BsaI, 400 U of T4 DNA ligase, 2 μL of T4 DNA ligase buffer (all from New England Biolabs, Ipswich, MA, USA), and deionized water to a final volume of 20 μL. The reaction was performed under the following conditions: 30 cycles of 37 °C for 5 min and 16 °C for 5 min, followed by 60 °C for 5 min and a 4 °C hold. PCR fragments used for Gibson assembly and Golden Gate assembly are listed in **Table 3**, and template plasmids, including MoClo Yeast Toolkit parts, are listed in **Table 1**. All constructed vectors were verified by Sanger sequencing (Macrogen, Seoul, Republic of Korea).

**Table 3.**
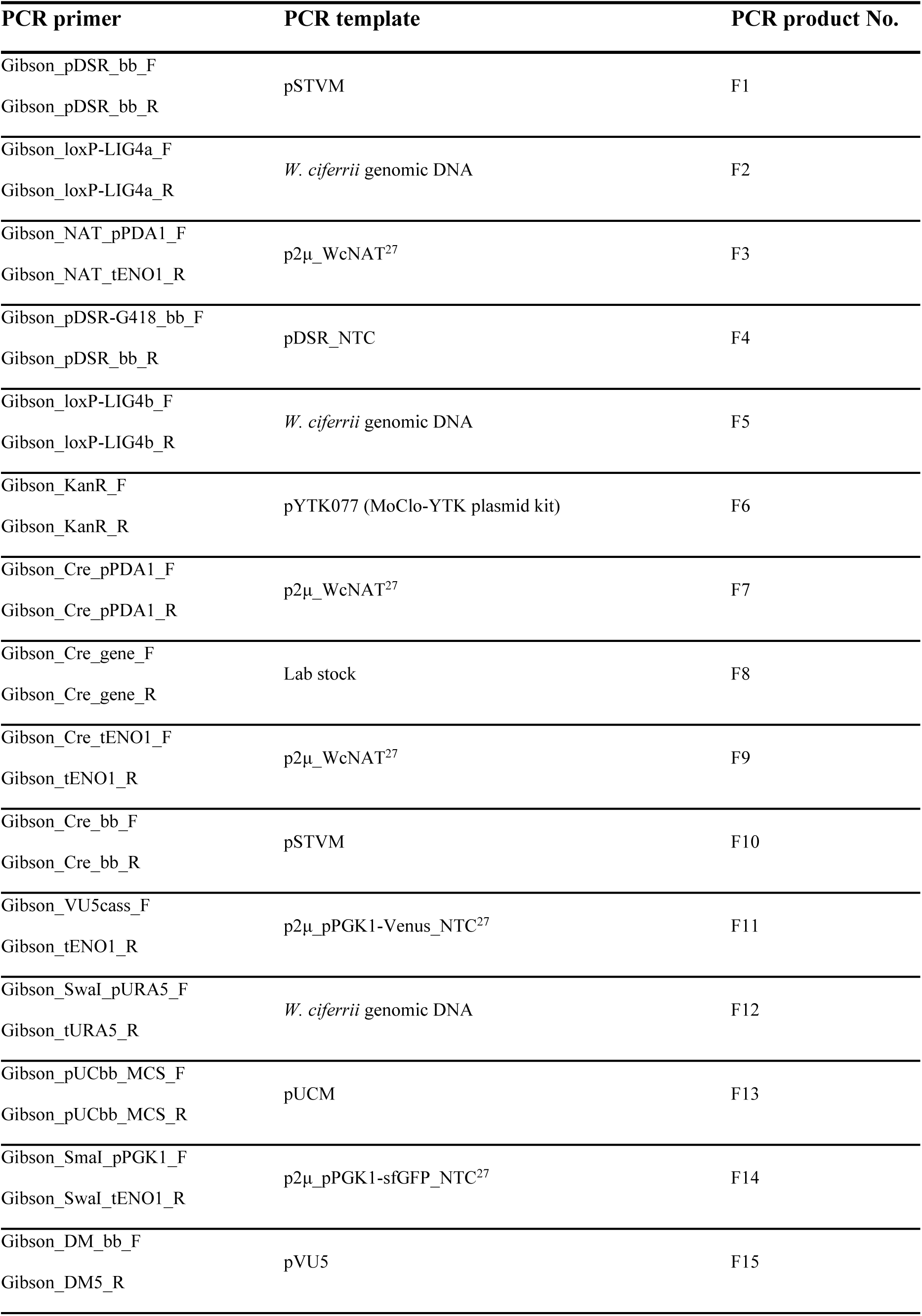
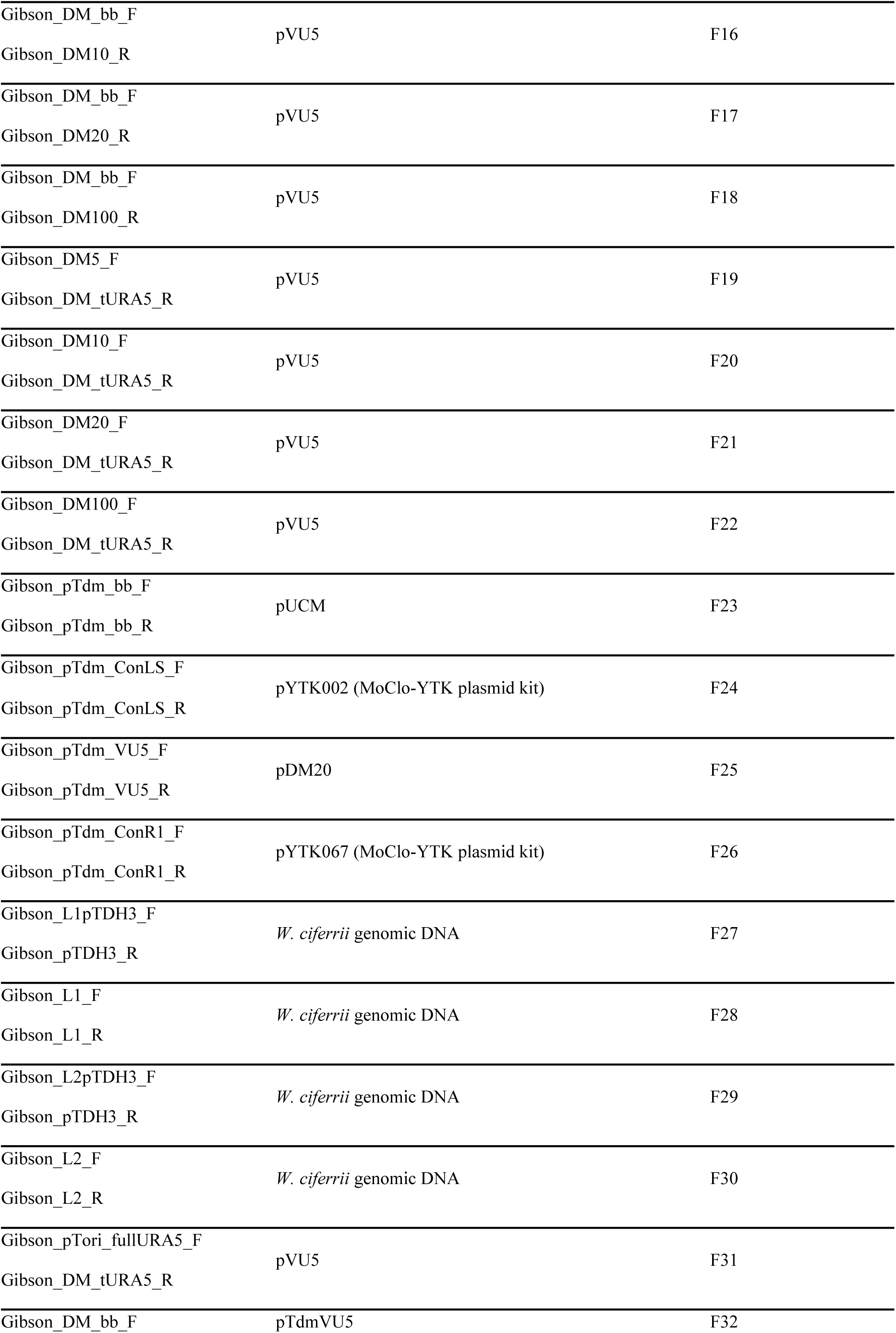
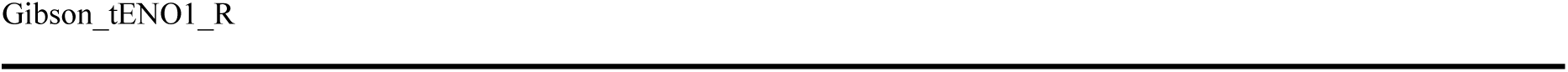
PCR fragment used for assembly in this study.

### Construction of *LIG4* disruption and Cre-recombinase vectors

To construct disruption vectors for biallelic deletion of *LIG4* in diploid *W. ciferrii*, two vectors carrying distinct antibiotic resistance markers were prepared. The *LIG4* gene sequence used for homologous recombination was identified from the *W. ciferrii* reference genome in the NCBI database (Gene ID: BN7_3078; RefSeq: NW_011887658.1) and confirmed by Sanger sequencing of genomic DNA extracted from the diploid production strain. The first disruption vector was designed for single-crossover homologous recombination and contained a loxP-flanked nourseothricin resistance cassette and a 1,013 bp internal fragment spanning positions 616–1628 of the *LIG4* coding sequence. The second vector was constructed analogously using a loxP-flanked geneticin resistance cassette and a 1,042 bp internal fragment spanning positions 216–1257. The nourseothricin and geneticin resistance cassettes were sourced from p2μ_WcNAT^27^ and pYTK077, respectively. Both vectors were assembled by Gibson assembly into the pSTVM backbone, which contains a chloramphenicol resistance marker and p15A origin of replication. Before transformation, the nourseothricin and geneticin disruption vectors were linearized by digestion with NdeI and PpuMI, respectively.

For Cre recombinase-mediated marker recycling, a transient Cre expression vector was constructed by cloning the bacteriophage P1-derived *Cre* coding sequence, 1,032 bp, under control of the endogenous *PDA1* promoter, 675 bp upstream of the start codon (Gene ID: BN7_3959a; RefSeq: NW_011887617.1), and *ENO1* terminator, 332 bp downstream of the stop codon (Gene ID: BN7_149; RefSeq: NW_011887739.1), into the pSTVM bacterial backbone by Gibson assembly. The resulting vector, pSTVM_Cre, was designed for transient expression and carried no yeast-compatible selectable marker or replication origin.

### Split-Venus reporter assay constructs

To assess DNA repair pathway preference in *W. ciferrii* and *S. cerevisiae*, split-Venus reporter templates were constructed by Golden Gate assembly. For the *W. ciferrii* assay, a Venus expression cassette was assembled into the p2μ_NTC backbone. The cassette consisted of the Venus coding sequence amplified from pYTK033 under control of the endogenous *PGK1* promoter, 497 bp upstream of the start codon (Gene ID: BN7_3735; RefSeq: NW_011887629.1), and *ENO1* terminator. For the *S. cerevisiae* assay, an analogous Venus expression cassette was generated using the *TEF2* promoter amplified from pYTK014 and the *ADH1* terminator amplified from pYTK053. In this construct, the antibiotic resistance marker was replaced with the *LEU2* auxotrophic marker amplified from pYTK075.

From each template vector, two PCR fragments sharing 50 bp homology arms within the Venus coding sequence were amplified. The overlapping regions corresponded to nucleotides 44–94 and 667–716 of Venus, and neither fragment alone encoded a functional fluorescent protein. Each fragment was gel-purified before transformation. For co-transformation, approximately 500 ng of each fragment was used for *W. ciferrii*, and approximately 200 ng of each fragment was used for *S. cerevisiae* CEN.PK2-1D.

### Construction of VU5 and telomeric repeat donor variants

For random integration screening using uracil auxotrophic selection combined with fluorescence-based readout, the pVU5 vector was constructed. A Venus expression cassette driven by endogenous *PGK1* promoter and *ENO1* terminator was linked to a functional *URA5* marker cassette containing the native *URA5* promoter, 500 bp upstream of the start codon, and *URA5* terminator, 141 bp downstream of the stop codon. The VU5 cassette was cloned into the pUCM backbone containing the ColE1 origin of replication and ampicillin resistance marker by Gibson assembly, yielding pVU5. For transformation, the VU5 cassette, 2,865 bp, was PCR-amplified from pVU5 using primers pPGK1_F and Gibson_DM_tURA5_R (**Table 2**) to generate linear donor DNA.

To generate telomeric repeat variants, PCR primers were designed to append 0, 1, 2, or 3 tandem copies of the 11 bp telomeric repeat sequence, 5′-CCCAGACACCA-3′, to both termini of the VU5 linear donor. Using pVU5 as the template, the 0× variant was amplified with primers pPGK1_F and Gibson_DM_tURA5_R, the 1× variant with primers 1-telo_F and 1-telo_R, the 2× variant with primers 2-telo_F and 2-telo_R, and the 3× variant with primers 3-telo_F and 3-telo_R (**Table 2**). The resulting PCR products were transformed into Δ*URA5* competent cells.

To determine whether the protective effect at the 22 bp optimum was specific to the telomeric sequence, two unrelated 22 bp sequences were used as length-matched negative controls: NC-1, 5′-GGACGCTTCAGCGATACTCCGC-3′, and NC-2, 5′-CCTGCGTCACGGTTGACCGTTC-3′. VU5 donor variants carrying these sequences at both termini were generated by PCR from pVU5 using primers NC1_F and NC1_R for the NC-1 variant, and NC2_F and NC2_R for the NC-2 variant (**Table 2**). The two control sequences were selected to match the 22 bp telomeric variant in length while sharing minimal local sequence identity with the telomeric repeat or with each other.

### Construction of GU5 reporter donors

To evaluate whether telomeric end-shielding was independent of the internal reporter sequence, the pGU5 vector was constructed by replacing the Venus cassette in pVU5 with a Superfolder GFP (sfGFP) cassette. Briefly, pVU5 was digested with SmaI and SwaI to remove the Venus cassette. The sfGFP cassette, consisting of pPGK1-sfGFP-tENO1, was PCR-amplified from the previously constructed p2μ_pPGK1-sfGFP_NTC reporter vector^27^. The two fragments were assembled by Gibson assembly to generate pGU5. GU5 linear fragments carrying 0× or 2× telomeric repeats were generated by PCR using primers pPGK1_F and Gibson_DM_tURA5_R for the 0× variant and 2-telo_F and 2-telo_R for the 2× variant (**Table 2**). The resulting donor fragments were transformed into Δ*URA5* competent cells.

### Construction of defective *URA5* marker donors

For the defective marker strategy, plasmids containing VU5 donor variants with progressively truncated *URA5* promoters were constructed. Using pVU5 as the template, linear PCR products were amplified with forward primers annealing 100, 20, 10, or 5 bp upstream of the *URA5* start codon and a reverse primer annealing to the 3′ end of the Venus *ENO1* terminator with a 5′ overhang complementary to the corresponding *URA5* promoter sequence. The resulting PCR products were circularized by Gibson assembly, yielding pDM100, pDM20, pDM10, and pDM5, named according to the length of the *URA5* promoter. Linear donor DNA fragments were then PCR-amplified from these plasmids using primers pPGK1_F and Gibson_DM_tURA5_R, and transformed into Δ*URA5* cells.

### Construction of pTdmVU5 and pToriVU5 platform vectors

The platform vector pTdmVU5 was constructed by Gibson assembly of four DNA fragments. The first fragment, corresponding to the bacterial backbone containing the ColE1 origin of replication and ampicillin resistance marker, was amplified from pUCM. The forward primer contained a 5′ overhang encoding the right multiple cloning site (MCS; NotI, SacII, ZraI, and AatII), and the reverse primer contained a 5′ overhang encoding the left MCS (XbaI, XmaI, and EcoRI), yielding a right MCS–backbone–left MCS fragment. The second fragment contained the left MoClo connector sequence, ConLS, amplified from pYTK002. The forward primer carried a 5′ overhang encoding the left MCS followed by two tandem copies of the 11 bp telomeric repeat, and the reverse primer carried a 5′ overhang encoding the SwaI recognition site and a loxP sequence. This generated a left MCS–2× telomeric repeat–ConLS–SwaI–loxP fragment. The third fragment, corresponding to the defective VU5 selection module, was amplified from pDM20. The forward primer shared 25 bp of homology with the terminal loxP sequence of the second fragment, and the reverse primer introduced a loxP site in the same orientation downstream of the *URA5* terminator. This fragment contained loxP(partial)–pPGK1–Venus–tENO1–20 bp *URA5* promoter–*URA5*–tURA5–loxP. The fourth fragment contained the right MoClo connector sequence, ConR1, amplified from pYTK067. The forward primer shared 25 bp of homology with the terminal loxP sequence of the third fragment, and the reverse primer carried a 5′ overhang encoding two tandem copies of the telomeric repeat followed by the right MCS sequence. The four gel-purified fragments were mixed at equimolar ratios and assembled using Gibson assembly.

The companion vector pToriVU5 was constructed identically to pTdmVU5 except that the 20 bp truncated *URA5* promoter in the defective VU5 module was replaced with the full-length 500 bp *URA5* promoter. For preliminary comparison of colony recovery under different selection stringencies, a heterologous donor fragment of approximately 8 kb, being characterized for an independent application, was inserted into both pTdmVU5 and pToriVU5 by Gibson assembly and transformed into Δ*URA5* competent cells.

### Construction of the *LCB1*–*LCB2* overexpression donor

For overexpression of *LCB1* and *LCB2*, pTdmVU5 was linearized with SwaI, and four PCR fragments were generated for Gibson assembly. The first fragment, containing the 415 bp *TDH3* promoter, was amplified from *W. ciferrii* genomic DNA using a forward primer with a 25 bp 5′ overhang homologous to the sequence immediately upstream of the SwaI cut site in pTdmVU5 and a reverse primer annealing to the promoter terminus, yielding a 443 bp product. The second fragment, spanning the *LCB1* coding sequence and its 309 bp terminator (Gene ID: BN7_169; RefSeq: NW_011887738.1), was amplified from *W. ciferrii* genomic DNA using a forward primer with a 25 bp 5′ overhang homologous to the 3′ end of the *TDH3* promoter and a reverse primer annealing to the terminator end, yielding a 2,047 bp product. The third fragment, corresponding to a second copy of the 415 bp *TDH3* promoter, was amplified using a forward primer with a 25 bp 5′ overhang homologous to the 3′ end of the *LCB1* terminator and a reverse primer annealing to the promoter terminus, yielding a 443 bp product. The fourth fragment, containing the *LCB2* coding sequence and its 277 bp terminator (Gene ID: BN7_5771; RefSeq: NW_011887508.1), was amplified from *W. ciferrii* genomic DNA using a forward primer with a 25 bp 5′ overhang homologous to the 3′ end of the second *TDH3* promoter and a reverse primer with a 25 bp 5′ overhang homologous to the sequence immediately downstream of the SwaI cut site in pTdmVU5, yielding a 2,016 bp product.

These four fragments were assembled into SwaI-linearized pTdmVU5 by Gibson assembly to generate pTdmVU5-pTDH3_*LCB1-*pTDH3_*LCB2*. Donor DNA was released from the assembled vector by XmaI–NotI double digestion, yielding an approximately 7.7 kb sticky-end fragment for transformation.

### Transformation of *W. ciferrii* and *E. coli*

Electroporation-based transformation of *W. ciferrii* was performed as described previously.^27^ Briefly, cells were grown in YPD medium to an OD_600_ of 0.8–1.2, harvested, resuspended in 0.01 culture volume of 50 mM phosphate buffer (pH 7.5), supplemented with 25 mM dithiothreitol, and incubated at 37 °C for 15 min. Cells were washed twice with one culture volume of STM solution containing 10 mM Tris-HCl (pH 7.5), 270 mM sucrose, and 1 mM MgCl_2_. The washed cells were resuspended in 0.01 culture volume of STM solution and stored at –80 °C in 50 μL aliquots.

For transformation, approximately 1 μg of DNA was mixed with 50 μL of thawed competent cells and electroporated at 500 V, 50 μF, and 700 Ω in a 0.2 cm cuvette using a GenePulser Xcell electroporator (Bio-Rad, Hercules, CA, USA). Cells were recovered in 2 mL of YPD medium at 25 °C for 6–12 h before plating onto selective agar. Transformation efficiency was calculated as colony-forming units per microgram of DNA. *E. coli* plasmids transformation was performed by electroporation into NEB 10-beta competent cells according to the manufacturer’s protocol.

### Fluorescence measurement

For fluorescence measurement, cells cultivated in 96-deep-well plates were transferred to 96-well black plates and analyzed using a Cytation 3 plate reader (BioTek, Winooski, VT, USA) in top-read mode. Venus and sfGFP fluorescence were both measured at excitation and emission wavelengths of 485 and 528 nm, respectively, with a gain setting of 50. OD_600_ was measured separately in 96-well clear plates using a SPECTRAmax PLUS384 spectrophotometer. Fluorescence values were normalized to OD_600_, and normalized fluorescence was further expressed relative to the non-fluorescent Δ*URA5* negative control included in each plate. When indicated, upper and lower outliers were excluded prior to statistical analysis using the same exclusion criterion across compared groups.

### Telomeric repeat sequence identification

The telomeric repeat unit of *W. ciferrii* was identified by tandem-repeat analysis of an in-house draft genome assembly currently under construction. Tandem Repeats Finder version 4.09 was run with the following parameters: match weight, 2; mismatch penalty, 7; indel penalty, 7; match probability, 80%; indel probability, 10%; minimum alignment score, 50; and maximum period size, 2,000 bp.^59^ The maximum period size was extended beyond the conventional 500 bp setting to capture short telomeric units and longer tandem-repeat arrays in a single analysis. The flanking sequence (-f), masked sequence (-m), and extended array-length (-l 6) output options were enabled. Repeat arrays located at one or both contig termini were inspected manually, and short tandem-repeat motifs recurring across multiple terminal positions were assigned as candidate telomeric repeats. Rotational permutations of the same motif were treated as a single repeat unit, and the canonical sequence was reported on the C-rich strand, consistent with fungal telomere convention.

### Copy number determination by qPCR

Genomic copy numbers of Venus, *LCB1*, and *LCB2* were determined by quantitative PCR using the endogenous *TDH3* gene as a diploid reference, assigned a copy number of 2. Cells were harvested by centrifugation, and genomic DNA was extracted as described above. qPCR reactions were performed using the SensiFAST™ SYBR No-ROX Kit (Meridian Bioscience, London, UK) on a Rotor-Gene system (Qiagen, Hilden, Germany). The cycling program was 95 °C for 2 min, followed by 40 cycles of 95 °C for 5 s, 60 °C for 10 s, and 72 °C for 5 s. The target gene and *TDH3* were amplified in parallel for each sample. Relative copy numbers were calculated using the ΔCt method, where ΔCt = Ct(target) – Ct(*TDH3*), and copy number was expressed as 2^−ΔCt^ × 2 to account for the diploid reference. All reactions were performed using three biological triplicates. Primers used for qPCR are listed in **Table 2**, designated as qPCR_gene_F and qPCR_gene_R.

### Time-course analysis of donor DNA persistence

To monitor donor DNA persistence after transformation, Δ*URA5* competent cells were electroporated with VU5 donor variants carrying 0, 1, 2, or 3 telomeric repeats and recovered in YPD medium at 25 °C with shaking at 250 rpm. At 1, 3, and 8 h post-transformation, 800 μL aliquots were withdrawn from the same recovery tube, washed three times with 1 mL of 1× phosphate-buffered saline (PBS) to reduce extracellular DNA carryover, and used for total DNA extraction as described above. Three independent transformation series were performed. qPCR was performed under the conditions described above using two primer pairs: qPCR_Venus_F/qPCR_Venus_R, targeting an internal region of the Venus coding sequence, and qPCR_pPGK1_F/qPCR_pPGK1_R, targeting the 5′ end of the *PGK1* promoter region of the donor DNA terminus. *TDH3* was used as the diploid reference gene, and relative copy numbers were calculated using the ΔCt method. The *PGK1* promoter 5′-end primer pair also amplified the endogenous *PGK1* a-allele, whereas sequence variation in the α-allele abolished primer binding. Therefore, the resulting copy-number values were corrected by subtracting one copy to remove the endogenous contribution and obtain the donor-derived terminal signal.

### Fluorescence-activated cell sorting (FACS)

Fluorescence-activated cell sorting was used to recover Venus-positive integrants that failed to form colonies on SD-Ura agar. Following transformation of the XmaI–NotI digested pTdmVU5-pTDH3_*LCB1*-pTDH3_*LCB2* donor DNA into Δ*URA5* competent cells, transformants were cultured overnight in YPD medium to allow outgrowth. A 1 mL aliquot of the outgrown culture was harvested by centrifugation and washed three times with 1 mL of 1× PBS. Cells were analyzed and sorted using a FACSAria III cell sorter (BD Biosciences, Franklin Lakes, NJ, USA) equipped with a FITC channel, with excitation at 495 nm and emission at 519 nm. The flow rate was maintained below 2,000 events per second. Venus-negative Δ*URA5* cells were used to define the background fluorescence gate, and cells showing a distinct fluorescence shift above the negative population were collected.

Approximately 10,000 Venus-positive events were sorted into YPD medium for recovery. Sorted cells were cultivated in liquid YPD medium at 25 °C until sufficient growth was observed. Cell density was estimated from OD_600_ using a conversion factor of approximately 3 × 10^7^ cells/mL per OD_600_ unit, and an appropriate volume was spread onto YPD agar to obtain isolated colonies. Ninety-six colonies were inoculated into a 96-deep-well plate containing 1 mL of YMglSC medium and cultivated at 25 °C and 800 rpm for 48 h.

Normalized fluorescence intensity was measured as described above, and the three clones exhibiting the highest values, designated 3F, 6C, and 12A, were selected for further characterization. During 96-deep-well plate cultivation, glycerol stocks were prepared in 96-well plate format at 24 h post-inoculation, when cells had reached logarithmic phase, enabling recovery of selected strains for flask-scale experiments.

### Flask-scale cultivation and TAPS production

For flask-scale TAPS production, wild-type *W. ciferrii*, the parental Δ*URA5* strain, and selected integrant strains were cultivated in 250 mL baffled flasks containing 50 mL of AAT medium. Cultures were incubated at 25 °C with shaking at 250 rpm. Biological triplicates were performed for each strain. Cell growth was monitored by measuring OD_600_, and culture samples were collected at the end of cultivation for TAPS extraction and quantification.

### Extraction and HPLC analysis of TAPS

For TAPS quantification, 1 mL of culture broth was withdrawn from each flask and mixed with 4 mL of methanol, followed by vigorous vortexing for 30 min. After extraction, the mixture was centrifuged, and the supernatant was filtered through a 0.2 μm PTFE syringe membrane filter. The filtrate was analyzed by high-performance liquid chromatography using an Agilent 1260 system (Agilent Technologies, Santa Clara, CA, USA) equipped with a photodiode array detector and a ZORBAX Eclipse XDB-C18 column (150 mm × 4.6 mm, 5 μm; Agilent Technologies) operated at 20 °C. Gradient elution was performed at a flow rate of 1 mL/min using mobile phase A, acetonitrile, and mobile phase B, deionized water. The gradient was as follows: 45–80% A from 0 to 18 min, 80–90% A from 18 to 20 min, 90–100% A from 20 to 23 min, 100% A from 23 to 25 min, 100–45% A from 25 to 28 min, and 45% A from 28 to 30 min. TAPS was detected at 200 nm and quantified based on peak area using an external calibration curve generated with a TAPS standard (Sigma-Aldrich, St. Louis, MO, USA).

### Transcriptional expression analysis

For transcriptional analysis, cells were harvested during logarithmic phase by centrifugation, and cell pellets were immediately resuspended in RNAlater stabilization solution (Thermo Fisher, Waltham, MA, USA) to preserve RNA integrity. Total RNA was extracted using the easy-BLUE™ Total RNA Extraction Kit (iNtRON Biotechnology, Seongnam, Republic of Korea) according to the manufacturer’s instructions. cDNA was synthesized using the ReverTraAce™ qPCR RT Kit (Toyobo, Osaka, Japan). Quantitative PCR was then performed using the SensiFAST™ No-ROX One-Step Kit (Meridian Bioscience, London, UK) on a Rotor-Gene cycler (Qiagen, Hilden, Germany) under the cycling conditions described above. Relative mRNA expression levels of *LCB1* and *LCB2* were calculated using the ΔΔCt method with *ACT1* as the reference gene and the parental Δ*URA5* strain as the normalization control. Primers are listed in **Table 2**. Each analysis was performed using three biological replicates.

### Statistical analysis

Data are presented as mean ± standard deviation (SD) unless otherwise stated. Statistical analyses were performed using GraphPad Prism version 8.0.2. Two-group comparisons were assessed using an unpaired *t*-test with Welch’s correction. For comparisons among three or more groups, Brown-Forsythe and Welch ANOVA followed by Dunnett’s T3 multiple comparisons test was performed relative to the designated control. Statistical significance was defined as *p* < 0.05. Significance labels are reported as follows: ns, *p* ≥ 0.05; \**p* < 0.05; \*\**p* < 0.01; \*\*\**p* < 0.001; \*\*\*\**p* < 0.0001.

## Conclusions

In this study, we developed a homology-independent multicopy genome integration platform for the non-model diploid yeast *W. ciferrii* by repurposing its native NHEJ activity. Rather than disrupting NHEJ to improve homologous recombination, which compromised cellular fitness and TAPS production, we exploited this dominant repair pathway as a mechanism for random genomic insertion. Through systematic donor design, we identified three key parameters that improve NHEJ-mediated integration: telomeric end-shielding to enhance linear donor persistence, defective *URA5* marker selection to enrich multicopy integrants, and 5′-phosphorylated donor termini to improve transformant recovery and reporter output.

These design principles were consolidated into the platform vector pTdmVU5 and applied to multicopy integration of *LCB1* and *LCB2*, which encode the two subunits of serine palmitoyltransferase. The resulting integrant strains showed increased gene copy number, elevated transcript abundance, and up to 2.7-fold higher TAPS production. We also identified a practical tradeoff between copy-number enrichment and donor-size tolerance, and developed a companion vector, pToriVU5, with the full-length *URA5* promoter as a complementary design for larger donor constructs.

Overall, this work establishes a practical strategy for rapid pathway amplification in *W. ciferrii* and provides a generalizable framework for exploiting NHEJ-dominant DNA repair in diploid non-model yeasts. By converting a major barrier to precise genome editing into a tool for multicopy integration, this platform expands the synthetic biology toolbox for industrial yeast strain development.

## Availability of data and materials

All data generated or analyzed during this study are included in this published article and its Supplementary Material.

## Acknowledgments

This work was supported by the National Research Foundation of Korea (NRF) (RS-2022-NR067491, RS-2025-23323220) and the Korea Institute of Marine Science and Technology Promotion (KIMST) funded by the Ministry of Oceans and Fisheries (RS-2022-KS221581).

## Author Contributions

PCL conceived the project. SRL and YS designed and performed the experiments. PCL, SRL, and YS wrote the manuscript. All authors edited and approved the manuscript.

## Declarations

### Ethics approval and consent to participate

This article does not contain any studies with human participants or animals performed by any of the authors.

### Consent for publication

Not applicable.

### Competing interests

The authors declare no conflict of interest.

## Notes

### Competing Interest Statement

The authors have declared no competing interest.

