## Supporting Information for "Repurposing native non-homologous end joining for multicopy random integration in *Wickerhamomyces ciferrii*"

*Corresponding Author

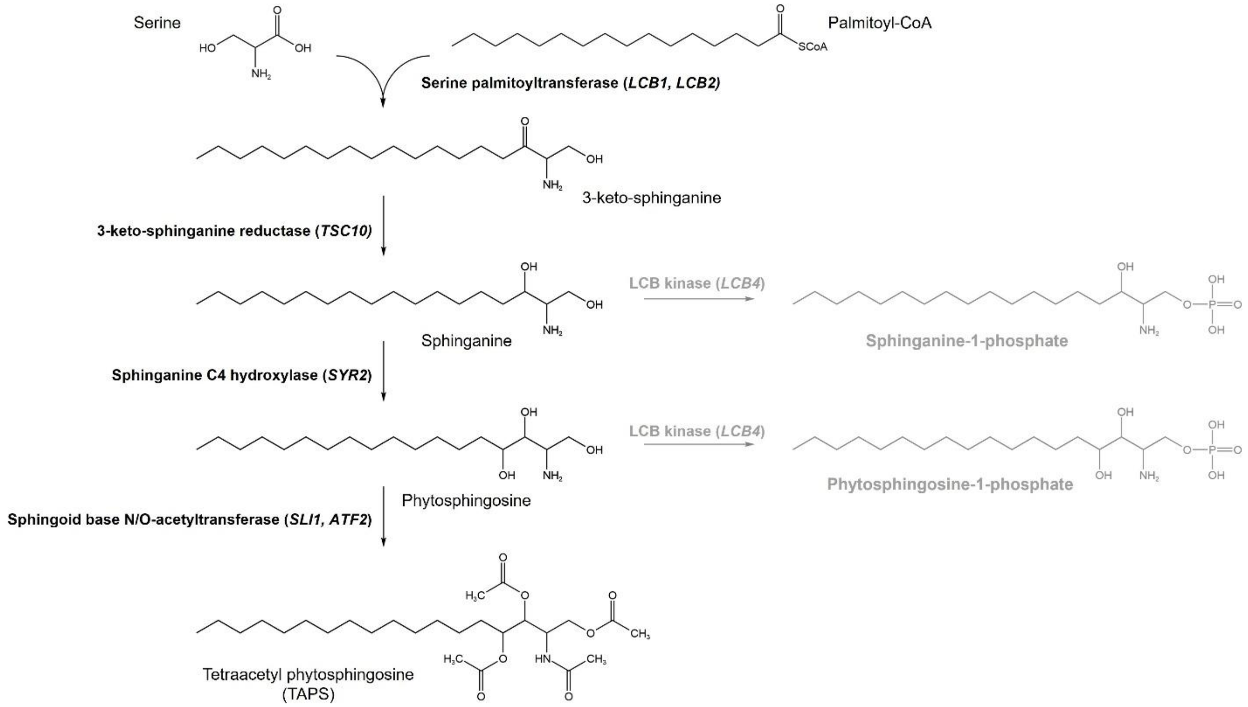


**Figure S1. Sphingolipid biosynthetic pathway leading to tetraacetyl phytosphingosine in *W. ciferrii*.** Serine and palmitoyl-CoA are condensed by serine palmitoyltransferase, composed of *LCB1* and *LCB2*, to form 3-keto-sphinganine. This intermediate is sequentially reduced by 3-keto-sphinganine reductase (*TSC10*), hydroxylated by sphinganine C4 hydroxylase (*SYR2*), and acetylated by sphingoid base N/O-acetyltransferase (*SLI1/ATF2*) to generate tetraacetyl phytosphingosine (TAPS). Competing phosphorylation reactions catalyzed by long-chain base kinase (*LCB4*) are indicated in gray.

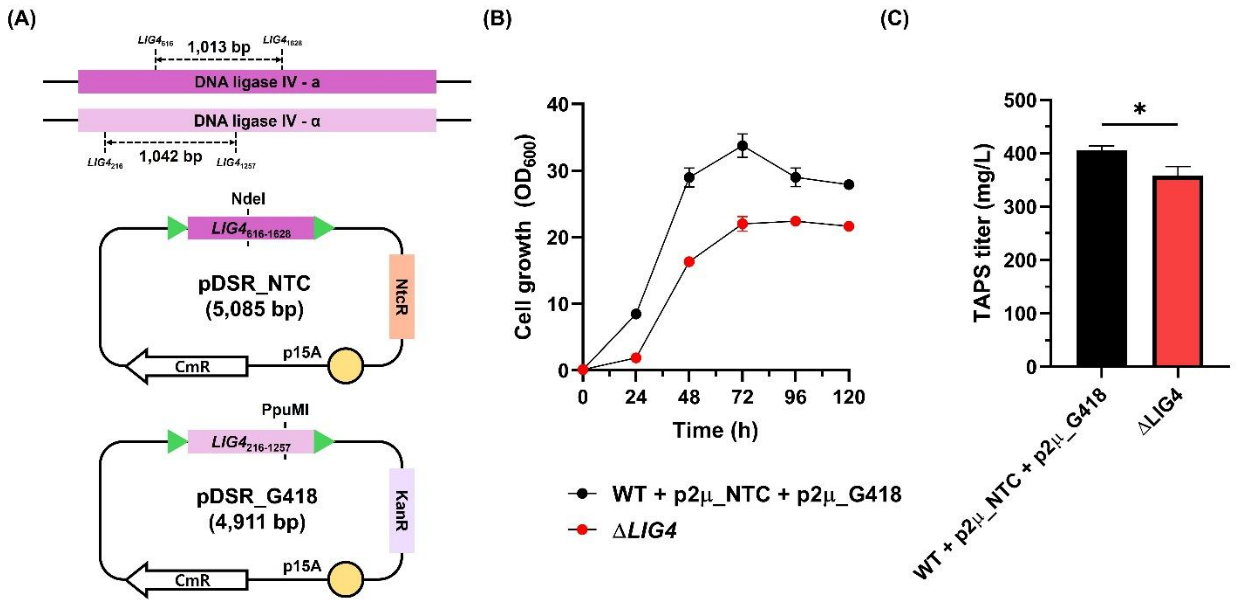


**Figure S2. Construction and phenotypic characterization of the biallelic Δ*LIG4* strain in diploid *W. ciferrii*.** **(A)** Schematic of the two single-crossover disruption vectors used for sequential disruption of the two *LIG4* alleles encoding DNA ligase IV. The pDSR_NTC vector carries a nourseothricin resistance marker and a 1,013 bp internal *LIG4* fragment, whereas pDSR_G418 carries a geneticin resistance marker and a 1,042 bp internal *LIG4* fragment. Restriction sites used for vector linearization are indicated. **(B)** Growth profiles of wild-type *W. ciferrii* carrying empty episomal plasmids, WT + p2μ_NTC + p2μ_G418, and the biallelic Δ*LIG4* strain cultured under antibiotic selection. **(C)** TAPS titers at the end of cultivation. Data represent the mean ± SD from three biological replicates. Statistical significance was determined using an unpaired *t*-test with Welch's correction (*$p < 0.05$).
